# Clinical Interventions and Inflammatory Signaling Shape the Transcriptional and Cellular Architecture of the Early Postnatal Lung

**DOI:** 10.1101/2025.10.17.683116

**Authors:** Tristan Frum, Angeline Wu, Marcela S. Ymayo, Yu-Hwai Tsai, Sha Huang, Mengkun Yang, Sydney G. Clark, Peggy P. Hsu, Ian A. Glass, Gail H. Deutsch, Jason R Spence

## Abstract

The early postnatal period in human development is characterized by extensive remodeling of the distal lung to support gas exchange, but this critical period remains poorly understood. Here, we constructed a comprehensive cellular atlas of the early postnatal human lung (0 to 2 years) using single-nucleus RNA sequencing of histologically normal specimens from 23 individuals. Our analysis identified two previously unknown and mutually exclusive transcriptional states of alveolar type 2 (AT2) cells, one defined by upregulation of genes involved in lipid metabolism and identified by unique expression of *FMO5*, and the other defined by upregulation of inflammatory response genes and identified by unique expression of *CFTR.* Using spatial transcriptomics, we discovered that AT2 cell states reside in specific niches and interact with distinct alveolar fibroblast subtypes. Clinical data and organoid experiments further suggest that the environment dictates which state prevails as the pro-inflammatory/pro-regenerative signals TNF-α and IL-1β promoted the *CFTR+* state *in vitro*, while patients treated with the anti-inflammatory drug dexamethasone (Dex) had more abundant *FMO5+* AT2 cells, and Dex induced the *FMO5+* state *in vitro*. These observations link inflammatory signaling and anti-inflammatory clinical interventions to shifts in transcriptional state of the alveolar epithelium. We benchmarked two neonatal lung diseases, bronchopulmonary dysplasia and pulmonary interstitial glycogenosis, revealing a profound disruption in the balance of AT2 states, a broad arrest of postnatal cellular development, and impaired cellular maturation. Our work uncovers a fundamental new understanding of early postnatal human lung biology, linking pro– and anti-inflammatory signaling to AT2 transcriptional phenotypes and providing a new framework for understanding lung disease.

## INTRODUCTION

During postnatal alveologenesis, the distal lung undergoes profound cellular and structural remodeling as it functionally matures in response to exposure to air^1–4^. In addition, the lung experiences dramatic growth during this period, and new alveolar units are formed^5–7^, while existing units undergo septation that drives expansion of the gas-exchange surface^8–10^. Lung injury during this period can be caused by premature birth, infection or other intrauterine/neonatal insults, leading to respiratory distress with possible permanent changes to lung function and maturation^11–14^. Studies in mice indicate that neonatal alveoli have enhanced growth and regenerative properties compared to adults^15–17^. Similarly, humans exhibit enhanced regenerative capacity during the neonatal period, where the lung can undergo compensatory growth after pneumectomy, capacity that is lost in older individuals^18–20^. Despite these unique properties that place the early postnatal lung at the nexus of alveolar development, functional maturation and regeneration, most single cell studies have focused on the fetal^21–25^ and adult lung^26–33^, with a few recent studies beginning to interrogate the neonatal and pediatric lung^34,35^. This lag likely reflects the difficulty of obtaining normal lung tissue from these early life stages.

To address this gap, we generated a single-nucleus RNA-sequencing (snRNA-seq) atlas of the early postnatal distal lung using flash-frozen histologically confirmed normal lung obtained from pediatric patients at autopsy (n = 3) and undergoing surgical resection for congenital airway malformations (n = 20) (Fig. 1A). This cohort covers the first two years of life (0-2 years), is sex-balanced and ancestrally diverse (Table S1). Through integrated analysis and iterative subclustering, we identified the full complement of expected epithelial, mesenchymal, immune, and endothelial populations as well as novel cell states within the alveolar epithelium. We also uncovered two previously uncharacterized states of AT2 cells: an *FMO5*+ state defined by a transcriptional program of hundreds of coexpressed genes highly enriched for lipid metabolic pathways, and a *CFTR*+ state comprising a similarly large gene set enriched for inflammatory pathway members. Using 10X Xenium spatial transcriptomics, we discovered that AT2 states are reflective of unique spatial niches composed of niche-specific alveolar fibroblast subtypes, AT2 to AT1 differentiating cells, and distinct signaling environments. We examined clinical data from our cohort, transcriptional differences in snRNA-seq and niche specific enrichment of ligands in spatial data to nominate and screen candidates in nascent AT2 organoids for the ability to drive *FMO5*+ or *CFTR*+ AT2 cell subtype identity and discovered that these AT2 cell states align along an axis of inflammatory and anti-inflammatory signaling. Notably, dexamethasone, a common clinical intervention in the context of neonatal lung injury, pushed the composition of AT2s in treated lungs towards an *FMO5*+ AT2 cell state *in vitro*, linking clinical interventions to shifts in AT2 transcriptome.

**Figure 1.**
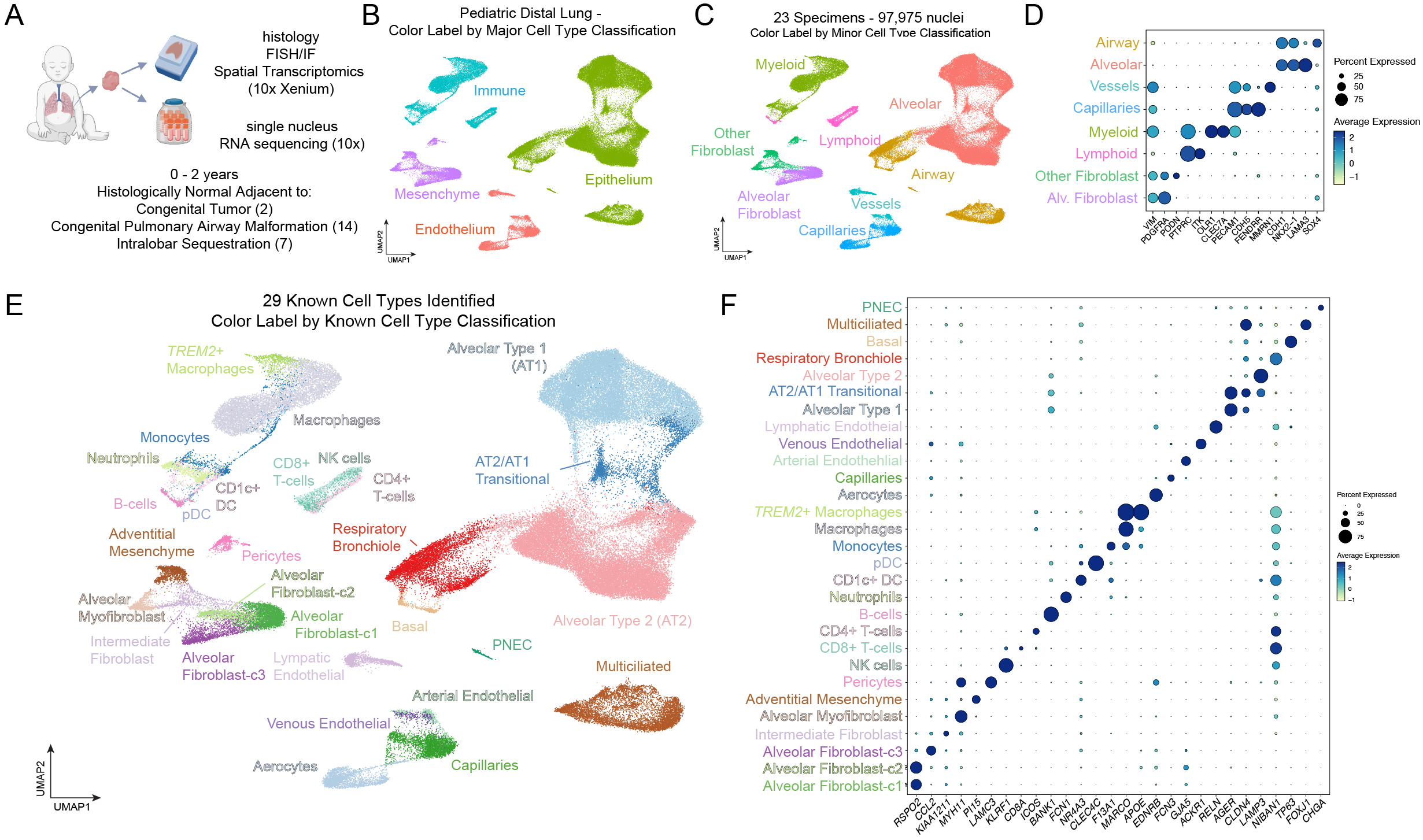
A Single-Cell Atlas of the Distal Human Lung from Birth to Two Years of Age. (A) Overview of study design. Flash-frozen distal lung tissue from 23 early postnatal patients (0–2 years) undergoing surgery for congenital malformations was analyzed by single-nucleus RNA-sequencing. Pathologist-identified histologically normal regions were selected for transcriptional profiling. (B) UMAP embedding of 97,975 nuclei after mutual nearest neighbor (MNN) integration across all specimens, showing separation of major cell classes. (C) Higher-resolution clustering of the dataset in (B) reveals clear segregation of airway and alveolar epithelial subtypes, lymphoid and myeloid immune cells, fibroblast subtypes (including Alveolar Fibroblast-c1/2), and endothelial versus capillary compartments. (D) Dot plot showing expression of key markers distinguishing major cell classes (*VIM*, *PTPRC*, *PECAM1*, *CDH1*) and representative markers of finer annotations (e.g., *LAMA3* for alveolar epithelium, *SOX4* for airway epithelium). (E) UMAP of snRNA-seq data annotated with known cell types assigned through iterative subclustering and curation of known markers (see Figure S1). (F) Dot plot showing expression of highly specific markers defining each annotated cell type in (E).

Additionally, we benchmarked two neonatal lung diseases, bronchopulmonary dysplasia (BPD) and pulmonary interstitial glycogenosis (P.I.G.) against our early postnatal atlas. This analysis revealed a profound disruption in the balance between mesenchymal and epithelial cells, a broad arrest of postnatal cellular development and a significant disturbance in the balance of *FMO5*+ and *CFTR*+ AT2 cell states, particularly in BPD.

Taken together, our findings uncover previously unrecognized facets of early postnatal lung biology with important implications for understanding the cellular and molecular underpinnings of human lung disease. These insights have the potential to guide clinical interventions, enhance disease monitoring through new biomarkers, and design new therapeutic strategies with awareness of the effect on AT2 cell state.

## RESULTS

### A sex-balanced and ancestrally diverse cell atlas of the early postnatal distal lung

To interrogate the early postnatal lung we performed snRNA-seq on flash-frozen distal lung specimens from patients ranging in age from within a week of birth to two years. Histologically confirmed normal lung was obtained from 3 patients who died from non-pulmonary causes and 20 individuals undergoing surgery for removal of localized congenital lung malformations or neoplasms (Fig. 1A). The patient cohort was from European (61%) and non-European ancestries (39%) and were sex-balanced (13 male, 10 female) (Table S1). Lung tissue was chosen for sequencing based on pathologist-selected normal regions from these specimens. Nuclei were isolated and sequenced using the 10X Chromium platform (see Methods). After ambient RNA removal and QC filtering, we obtained 97,975 high-quality nuclear transcriptomes. Integration of transcriptomic data from all specimens using a mutual nearest neighbor approach (*FastMNN*) led to clear segregation of major cell classifications (epithelium, mesenchyme, immune, endothelial) (Fig. 1B) and further delineation into more refined subtype classifications (e.g. alveolar, airway) (Fig. 1C-D). To identify specific cell types in our data each major cell classification was extracted and subclustered at a resolution that was sufficient to resolve known cell types by well-established cell type-specific transcriptional markers, identifying 29 known cell types in our dataset (Fig. 1E-F).

In the mesenchyme (Fig. S1A) we identified three clusters of alveolar fibroblast (i.e. lipofibroblast) cells coexpressing the canonical marker *ITGA8*^36^ along with *PDGFRA*, *PIEZO2*, *NPNT*, including one population (alveolar fibroblast-c3) that was defined by *CCL2* expression (Fig. S1B). Non-alveolar fibroblast cell clusters were distinguished from alveolar fibroblast cell clusters by coexpression of *ZEB1* and *SVIL* and consisted of alveolar myofibroblasts (*MHY11*/*ACTA2+)*, adventitial fibroblasts (*PI15*/*SCARA5+*), and pericytes (*PDGFRB*/*MYO1B+)* (Fig. S1B). There was also a population of ‘intermediate’ fibroblasts located centrally on the UMAP coexpressing markers of both alveolar fibroblast cells and other mesenchymal cell types (e.g. *PDGFRA*/*ZEB1*+) (Fig. S1A-B).

In the endothelium (Fig. S1C), capillaries were distinguished from populations in larger vessels by the expression of *ADGFRL2* and *FENDRR*, expressed the lung specific capillary marker *CA4*^37,38^ and contained robust aerocyte^27,38,39^ (aCAP; *HPGD*/*EDNRB2*+) and general capillary (gCAP, *BMP6*/*IL7R*+) populations. Arterial (*DKK2*/*GJA5*+), venous (*ACKR1*/*VCAM1*+), and lymphatic (*CCL21*/*RELN+*) cells were readily distinguished in the remaining endothelial clusters (Fig. S1D).

In the immune compartment (Fig. S1E) lymphoid cells were readily distinguished from myeloid based on *ETS1* and *CD96* expression and included natural killer (NK-) (*KLRF1*/*LINGO2*+), CD4+ T-(*CD28*/*ICOS*+), CD8+ T-(*ITGA1*/*CD8A*+), and B-cells (*BANK1*/*MS4A1*+). Myeloid cells were distinguished by broad *CD74*, *PLXDC2* and *SLC11A1*, expression and included neutrophils (*CSF3R*/*FCN1*+), CD1c+ DCs (*NR4A3*/*GPAT3*/*CD1C+*), plasmacytoid DCs (*RUNX2*/*CLEC4C+*), monocytes (*FMN1*/*RGL1+*), abundant macrophages (*MARCO*+) (Fig. S1F), and a macrophage subset expressing markers of ‘foamy’ or ‘lipid laden’ macrophages (*APOE*/*NPC2*/*TREM2*+) which was also enriched in our specimens for genes encoding complement, cathepsins, lysozyme, and S100 protein family members (Fig. S1I). We refer to this cluster as TREM2+ macrophages, as this is a well-studied macrophage subset marker indicative of a polarized, anti-inflammatory and lipogenic state^40–43^.

In the epithelium (Fig. S1G), alveolar type 1 (AT1, *CAV1*/*RTKN2*/*AGER*), and AT2 cells (*SFTPC*/*LAMP3*/*ETV5*+) were distinguishable, along with cells located between AT1 and AT2 cells on the UMAP embedding. These cells coexpressed markers of AT1 and AT2 cells and likely correspond to cells undergoing AT2 to AT1 differentiation (Fig. S1H). We also detected cell populations localized in distal respiratory bronchioles, which broadly expressed *SFTPB, SCGB3A2*, and *SOX4* (Fig. 1D, Fig. S1H). These included TRB/RASCs (*CP*/*RNASE1*+)^31,32^, pulmonary neuroendocrine cells (*CHGA*+) consisting of both *GRP*+ and *GHRL*+ subtypes^22,44^ (Fig. S1H,J), basal cells (*TP63/KRT17/KRT5*+) expressing the distal basal subtype marker *SFTPB*^31^ and multi-ciliated cells (*CFAP733/FOXJ1*+) exhibiting a *C6*+/*MUC16*-signature^45^ (Fig. S1H,K).

To interrogate changes in cell composition that coincide with dramatic changes in appearance of the distal lung during the transition from fetal to early postnatal periods of life (Fig. S1L) we compared our early postnatal atlas to available distal fetal lung snRNA-seq data (14 – 16 weeks)^24^. This comparison showed a dramatic shift from mesenchymal to epithelial cell dominance in early postnatal specimens (Fig. S1M). Myeloid cells were the dominant immune cell type and comprised similar proportions of the immune compartment in fetal and early postnatal specimens (Fig. S1N). Representation of the immune system increased proportionally and showed signs of increasing complexity, such as the onset of *MARCO* expression in the monocyte lineage, indicative of emergence of tissue resident macrophages^46–49^ (Fig. S1O). Maturation was also apparent in the capillary lineage, with early postnatal specimens showing clear increases in aerocyte (aCAP) and general capillary (gCAP) transcriptional signatures defined in adult specimens^27^ (Fig. S1P).

### Identification of transcriptionally heterogenous AT2 states in the alveolar epithelium during early life

In order to more deeply interrogate AT2 and AT1 cells, we computationally extracted and re-clustered epithelial cell types specific to the alveolus. Louvain clustering of epithelial cells at high resolution revealed multiple AT2, AT1 and AT2 to AT1 transitional cell subclusters within the data (Fig. 2A). From the standpoint of marker gene expression, it appeared that AT1 subclusters and AT2 to AT1 transitional subclusters represented cellular transitional states along an AT2 to AT1 differentiation trajectory (Fig. 2B). In addition, gene set enrichment analysis of AT1 and AT2 to AT1 transitional clusters revealed enrichment of terms related to cellular activities previously established to be involved in AT1 differentiation, including G protein-coupled receptor signaling^50^, semaphorin signaling^34,51^ cell adhesion and extracellular matrix organization^52^ (Fig. S2A). We identified two additional clusters of AT2 cells expressing equal levels of canonical AT2 marker genes (*SFTPC, ABCA3*, *LAMP3 etc.)* with no expression of AT2 to AT1 transitional or AT1 markers (Fig. 2B). GSEA of the top 200 genes enriched in either AT2 cluster (Fig. S2B) revealed that AT2-cluster 1 (AT2-c1) and AT2-cluster 2 (AT2-c2) were distinguished by enrichment of genes encoding important enzymes and transporters involved in lipid metabolism (AT2-c1: *CD36, HMGCS1, PLA2G4F, FASN*) (Fig. 2D, Fig. S2C), or inflammation related genes known to be activated by TNF-α or NF-κB activity (AT2-c2; *NFKBIA, RELB, NAMPT*) (Fig 2D, Fig. S2D,G).

**Figure 2.**
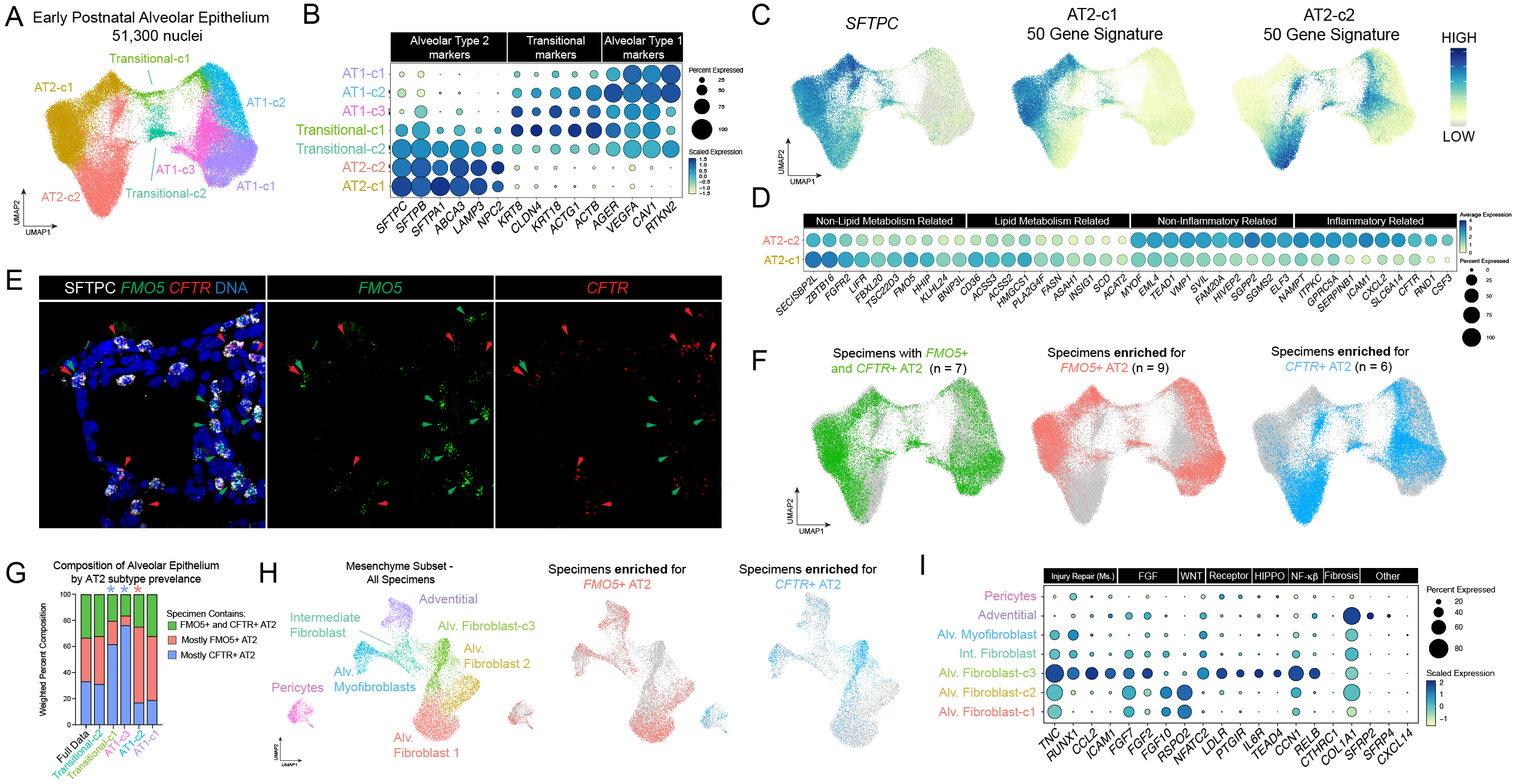
Transcriptionally Heterogenous AT2 cells Associate with Distinct Alveolar Fibroblast Subsets. (A) UMAP embedding of the early postnatal alveolar epithelium (51,300 nuclei) showing high-resolution subclustering of AT2 cells, AT2 to AT1 transitional cells, and AT1 cells. (B) Dot plot showing expression of marker genes defining AT2, AT2 to AT1 transitional, and AT1 states. The gradual loss of AT2 markers and acquisition of AT1 markers places transitional subclusters along an AT2-to-AT1 differentiation trajectory but does not account for transcriptional heterogeneity among AT2 subclusters. (C) Feature plots showing mutually exclusive module-score expression of the top 50 genes enriched in AT2-c1 and AT2-c2. (D) Dot plot summarizing differential expression across AT2 subclusters. AT2-c1 is enriched for genes involved in lipid metabolism, whereas AT2-c2 is enriched for genes associated with inflammatory and immune signaling. Both subclusters also display additional pathway enrichments not directly related to these categories, indicating broader transcriptional divergence between lipogenic and immune-responsive AT2 states. (E) Fluorescence *in situ* hybridization for *FMO5* and *CFTR* combined with immunostaining for the AT2 marker SFTPC confirms spatial separation of AT2 subtypes in situ. Red arrows indicate *CFTR*-high/*FMO5*-low cells, green arrows indicate *FMO5*-high/*CFTR*-low cells, and yellow arrows denote cells co-expressing both genes. (F) UMAPs highlighting specimen-specific enrichment of AT2 subtypes. Cells from each specimen classification are colored, with all other cells shown in gray. (G) Weighted composition of alveolar epithelial cell types across specimen categories based on AT2 subtype prevalence. AT2 to AT1 transitional 1 and AT1-c3 cells are over-represented in specimens enriched for AT2-c2 (blue stars), whereas AT1-c1 and AT1-c2 cells are over-represented in specimens enriched for AT2-c1 (red stars). (H) UMAP of subclustered mesenchymal nuclei showing the relationship between epithelial subtype enrichment and fibroblast composition. Specimens enriched for AT2-c1 (red) contain predominantly alveolar fibroblast-c1 and alveolar fibroblast-c2 populations, while specimens enriched for AT2-c2 (blue) contain mostly alveolar fibroblast-c3 cells. (I) Dot plot showing differential gene expression among alveolar fibroblast subclusters, highlighting distinct pathway enrichments related to FGF, WNT, HIPPO, NF-κB, and fibrosis-associated signaling.

To confirm transcriptional heterogeneity within early postnatal AT2s we selected highly enriched markers of each AT2 cluster (AT2-c1: *FMO5,* AT2-c2: *CFTR*) (Fig. S2E-F) and performed fluorescence *in situ* hybridization (FISH) on matched sections from our sequenced cohort for *FMO5* and *CFTR* in combination with immunofluorescence for SFTPC. Using this approach, we readily distinguished SFTPC+ AT2s with enriched *FMO5* expression from those with enriched *CFTR*, with a small minority of AT2s expressing both markers (Fig. 2E).

Based on the enrichment of genes involved in lipid metabolism in one subtype, marked in tissue by the distinct expression of *FMO5* (AT2-c1), and enrichment of inflammation-related transcripts in the other subtype, marked in tissue by the expression of *CFTR* (AT2-c2), we refer to these subclusters as “*FMO5*+ AT2s” and “*CFTR*+ AT2s” throughout the remainder of the manuscript.

In the adult human lung, unique transcriptional signatures of AT2s have been described including a ‘canonical’ AT2 signature and a less widely expressed ‘signaling’ AT2 (AT2-s) signature defined by enrichment for WNT pathway components^27^. To interrogate how early postnatal AT2 signatures identified in our data align with these previously defined adult AT2 subtype signatures we examined the expression of previously defined markers of canonical AT2s and AT2-s in *FMO5*+ and *CFTR*+ early postnatal AT2s (Fig. S2H) and performed gene module scoring on both early postnatal and adult AT2 subclusters using modules comprised of the top 50 genes differentially expressed between *FMO5+* and *CFTR+* AT2 subclusters (Table S2) or canonical AT2 and AT2-s cells (Table S3) (Fig. S2I). Early postnatal AT2 subclusters did not show differential expression of markers noted previously to be differentially expressed between adult AT2 subtypes (Fig. S2H); however using a broader ‘canonical’ AT2 marker set, *CFTR*+ AT2s scored higher for the ‘canonical AT2’ signature (Fig. S2I). In contrast both early postnatal AT2 subclusters scored low for the AT2-s signature (Fig. S2I, left panels). Likewise, neither adult AT2 subcluster showed differential enrichment for the transcriptional signatures defining *FMO5*+ and *CFTR*+ AT2s in the early postnatal lung (Fig. S2I, right panels). These findings suggest that *CFTR*+ AT2s are more similar to adult canonical AT2s than *FMO5*+ AT2s, and that the *FMO5*+ to *CFTR*+ transcriptional axis is distinct from the canonical AT2 to AT2-s axis in adult lungs.

### *FMO5*+ AT2 and *CFTR*+ AT2s associate with distinct alveolar fibroblast subtypes

The distribution of *FMO5*+ and *CFTR*+ AT2 states across individual specimens revealed three patterns of AT2 subtype abundance: (1) specimens highly enriched for lipogenic *FMO5*+ AT2s (n = 9); (2) specimens highly enriched for pro-inflammatory/pro-regenerative *CFTR*+ AT2s (n = 6); (3) specimens containing relatively equal proportions of both AT2 transcriptional states (n = 7) (Fig. 2F). One specimen was not classified due to low representation of AT2 cells (Table S1). We used these classifications as the basis for a stratified analysis of cell type abundance across the major cellular compartments (epithelial, mesenchymal, immune, and endothelial) in each AT2 subtype abundance category. For each group (i.e., *FMO5+* AT2-enriched, *CFTR+* AT2-enriched, and mixed specimens; Fig. 2F), cells were weighted so that each group contributed equally to the overall dataset (first column, Fig. 2G; Fig. S2L,N,P,R). We then used the weighted contributions of individual cell types within each compartment to identify associations of other cell types with the presence of *FMO5*+ or *CFTR*+ AT2s. In the alveolar epithelium, specimens enriched for *CFTR+* AT2s showed a higher relative contribution of AT2 to AT1 transitional cells and AT1-c3 cells (Fig. 2G). In contrast, specimens enriched for *FMO5+* AT2s (AT2-c1) contributed disproportionately more to the AT1-c2 subclusters (Fig. 2G). Enrichment of *CFTR+* AT2s in a specimen also correlated with the presence of AF-c3 cells in the mesenchyme (Fig. 2H, Fig. S2N). AF-c3 cells uniquely expressed markers previously associated with regeneration in the alveoli of mice^53–55^ (*TNC, RUNX1, CCL2, ICAM1*; Fig. 2I) and displayed altered profiles of FGF-, WNT– and other signaling ligands (Fig. 2I). In contrast, they were negative or low for markers of fibrosis-associated fibroblast populations like *CTHRC1* and *COL1A1* (Fig. 2I). A *CCL2*+ fibroblast population appears during regeneration in the mouse lung; however, human AF-c3s were negative for other markers defining this population in mice, such as *SFRP2*, *SFRP4* and *CXCL14*^54^ (Fig. 2I). Specimens with *FMO5+* AT2s and specimens with both *FMO5+* and *CFTR+* AT2s predominantly possessed AF-c1 and AF-c2 cells. In contrast to AF subcluster differences, specific endothelial and immune cell types did not correlate with AT2 subtype (Fig. S2O–R).

These observations led us to hypothesize that the presence of *CFTR*+ and *FMO5*+ AT2s reflects distinct stem cell niches composed of different AF populations and AT2 to AT1 transitional intermediates. To test this hypothesis, we performed 10X Xenium spatial transcriptomics on n = 5 specimens (Fig. 3A) using a customized 480 target gene panel designed to capture the full cellular heterogeneity in the early postnatal lung as based on our snRNA-seq data (Fig. S3A, Table S4). When combined with the Xenium *In Situ* Multimodal Cell Segmentation kit can overlay precise cell identity calls onto imaging data. Cells were segmented and transcripts assigned to cells using Xenium Ranger to generate a single-cell expression matrix. Cell identities were assigned by computational label transfer from our annotated early postnatal snRNA-seq dataset onto the Xenium data (Fig. 3B). AF-c1 and AF-c2 were collapsed to a single annotation for this analysis due to their similarity and common association with *FMO5*+ AT2s (hereafter AF-c1/2). We then used Seurat’s built-in ‘niche’ analysis, which considers the local composition of cell types to define areas of distinct cellular composition across tissue through a k-means based classification. To align niches across tissue sections we performed Pearson’s correlation using the cell type composition for each identified niche. (Fig. 3C) Using this approach, we distinguished bronchiolar and interstitial regions within tissue sections (Fig. 3C-D, light blue and purple regions). In addition, we identified two distinct cellular niches within the alveolar areas of tissue sections (Fig. 3C-D, red and gold regions). In some instances, alveolar niches were intermixed in a ‘salt and pepper’ pattern (Fig. 3D, top panel), while in other cases, niches were localized distinctly and associated with differences in the architecture of alveoli (Fig. 3D, bottom panel). Comparison of the cell type composition of the two alveolar niches revealed that alveolar niche 1 was significantly enriched for *FMO5*+ AT2s, AT1-c2, and AF-c1/2, whereas alveolar niche 2 was significantly enriched for *CFTR*+ AT2s, AT1-c3, and AF-c3 (Fig. 3E). Therefore, spatial transcriptomics analysis aligned well with the inference of cell type associations from stratified analysis of specimens (Fig. 2F-H, Fig. S2K-R) and confirmed that associations of AT2 subclusters and their cognate AF subcluster occur at a local level.

**Figure 3.**
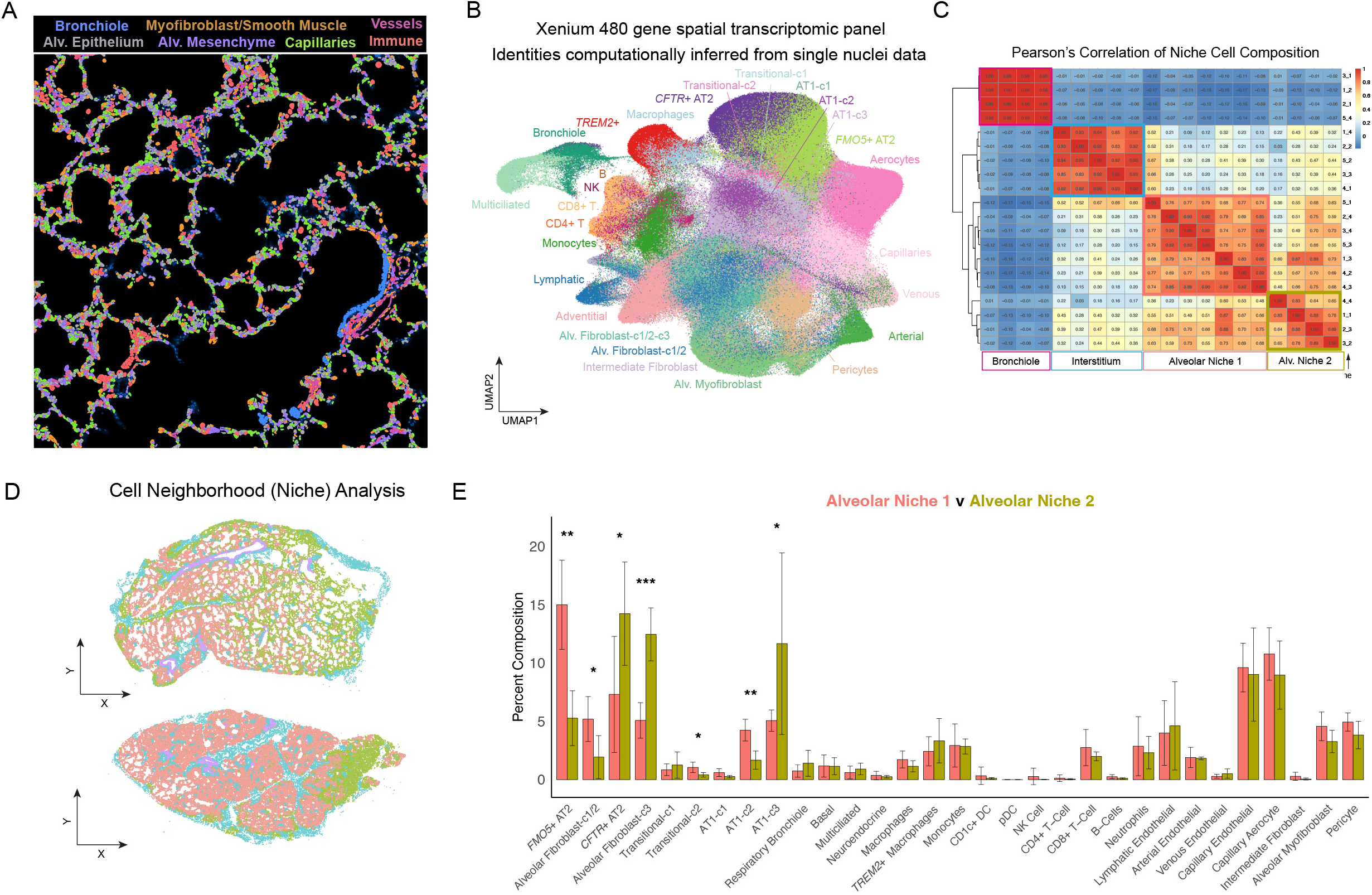
Spatial Association of *FMO5*+ and *CFTR*+ AT2s with Alveolar Fibroblast Subsets Defines Two Alveolar Niches in the Early Postnatal Lung. (A) Segmented spatial transcriptomic map of the early postnatal distal lung generated using the 480-gene Xenium panel, visualized in Xenium Explorer. Major tissue compartments, including bronchioles, alveolar epithelium, mesenchyme, vessels, and immune cells, are shown. (B) Integrated UMAP of all segmented cells from spatial transcriptomic sections, with cell identities computationally inferred by label transfer from snRNA-seq data. (C) Heatmap showing Pearson’s correlation of niche cell composition derived from Seurat v5’s k-means–based niche analysis across all five specimens. Hierarchical clustering identifies recurring spatial neighborhoods, including respiratory bronchioles, interstitial regions, and two distinct alveolar niches. (D) Spatial maps of two early postnatal lung sections, colored by niche classification, illustrating the distribution of alveolar, interstitial, and bronchiolar regions across the tissue. (E) Bar plot comparing proportional cell type composition between Alveolar Niche 1 (red) and Alveolar Niche 2 (gold). Niche 1 is enriched for *FMO5*+ AT2 cells, alveolar fibroblast-c1/2 populations, and AT2 to AT1 transitional-c2 and AT1-c2. Niche 2 is enriched for *CFTR*+ AT2 cells, alveolar fibroblast-c3 cells, and AT1-c3. Statistical significance was determined by Welch’s t-test using each section as a biological replicate (*p < 0.05; **p < 0.01; ***p < 0.001).

To validate this finding in a supervised manner we also annotated by hand the AT2 and mesenchymal cells in our spatial data and performed additional analysis to evaluate the association between AT2 states and AF subtypes. Using markers of *FMO5*+ and *CFTR*+ AT2s included in our Xenium panel, AT2s were delineated into three categories: (1) *CFTR*+ AT2s expressing *CFTR*+ signature genes and negative for *FMO5*+ signature genes; (2) *FMO5*+ AT2s expressing *FMO5*+ signature genes and negative for *CFTR*+ signature genes; and (3) AT2s expressing neither, termed ‘Unassigned’ (Fig. S3B). Mesenchymal populations were also well distinguished in Xenium data, with AF-c3 readily distinguished from other fibroblast populations based on *CCL2* expression (Fig. S3C). Assigning AT2 and AF identities back to the spatial data and ignoring all other cell types (Fig. S3D) we then measured the average distance between *FMO5+* AT2s, *CFTR+* AT2s, AF-c1/2 cells 1/2 and AF-c3 cells (Fig. S3E). This analysis showed that *FMO5+* AT2s were closest with AF-c1/2, *CFTR+* AT2s were closest to AF-c3, and *FMO5+* AT2s were furthest from AF-c3, indicating they are anti-correlated in space (Fig. S3E). We also performed niche analysis using just *FMO5+* AT2s, *CFTR+* AT2s, AF-c1/2, and AF-c3 cells, which identified two alveolar niches defined by the association of *FMO5*+ AT2s with AF-c1/2 cells, and *CFTR*+ AT2s with AF-c3 cells, concurring with our analysis using cell type identities inferred by label transfer (Fig. S3F-H). Finally, to examine the self-association of AT2 subtypes we performed join count-based clustering of AT2 subtypes and found that *CFTR*+ AT2s were more likely to be in contact with each other than *FMO5*+ AT2s (Fig. S3J). Taken together, our data identify two spatially distinct AT2 niches that correspond to specific fibroblast populations in early postnatal lungs.

### AT2 transcriptional signatures are regulated along an inflammatory axis

Having determined the close spatial association of AT2 subtypes and their cognate AF subtypes, we analyzed our snRNA-seq data using CellChat^56,57^ to model how cell signaling interactions would differ between the alveolar epithelium and mesenchyme in the two alveolar niches defined by our spatial analysis (Fig. 3, Fig. S3). Focusing first specifically on communication between AF subclusters and their cognate AT2 subclusters (i.e. *FMO5+* AT2 with AF-c1/2, *CFTR+* AT2 with AF-c3) (Fig. S4A-F) we determined that BMP, EGF, and FGF signaling, known regulators of the AT2 cell lineage, were differentially active between the two niches (Fig. S4C-D). At the level of ligands and their sources these pathway level changes encompassed decreased *BMP5* and increased *FGF2* and *SEMA3C* in AF-c3’s relative to AF-c1/2 (Fig. S4E), and a gain in *BMP2* from *CFTR+* AT2s relative to *FMO5+* AT2s (Fig. S4F). *CFTR*+ AT2s also uniquely express *AREG* and *CSF3*, which have the potential to act in an autocrine manner (Fig. S4F).

To look more broadly and include endothelial and immune cell types in our analysis we relied on our stratification of specimens by AT2 subtype abundance (Fig. S2K-Q) and compared cell signaling between *FMO5*+ AT2 and *CFTR*+ AT2 enriched specimens (i.e. *FMO5+* AT2 specimens n = 9; *CFTR+* specimens n = 6) (Fig. S4G-H). This analysis identified the increase of several macrophage-derived cytokines in *CFTR+* AT2 specimens including *IL-6*, *TNF-*α, *IL-1*β, and, as expected, *CCL2* (Fig. S4G-H). We confirmed these genes were localized to the *CFTR*+ AT2/AF-c3-enriched alveolar niche 2 in our spatial data (Fig. S4I).

Dexamethasone (Dex) or other glucocorticoids are often used clinically as anti-inflammatory agents, and to help enhance lung maturation *in utero* in pregnant women at risk of premature birth^57^. Within our sequenced specimens, several patients had recently received Dex or other glucocorticoid steroid treatment (n = 10). We therefore stratified all samples as ‘Dex treated’ or ‘Dex naïve’ and quantified the number of *FMO5+* or *CFTR+* AT2s, finding that >75% of AT2 cells in Dex treated patients were *FMO5+* compared to ∼35% in Dex naïve patients (Fig. 4A). To directly test if Dex, among other pathways, influence the AT2 lineage toward an *FMO5+* or *CFTR+* transcriptional signature we utilized a dexamethasone-free AT2 differentiation method to generate nascent AT2 organoids^58^ from lung bud tip progenitor organoids^21,59,60^ and nominated thirty-one growth factors, cytokines, small molecules or metabolites selected based on clinical information (i.e. Dex), CellChat analysis or differential expression analysis (upregulated genes/pathways) (Table S5) and performed a screen for inducers of the *FMO5+* or *CFTR+* phenotype in nascent AT2 organoids (Fig. 4B-C). This screen included 8 genes expressed in the *CFTR+* AT2 signature and 7 genes expressed in the *FMO5+* AT2 signature and was replicated in three independent biological specimens (Fig. 4C). We identified the cytokines TNF-α and IL-1β as promoting a *CFTR+*-like AT2 transcriptional phenotype and confirmed Dex as promoting a *FMO5*+-like phenotype (Fig. 4C). We further confirmed a similar response to TNF-α, IL-1β and Dex using primary AT2 organoids derived from an early postnatal lung specimen (Fig. S4J-K).

**Figure 4.**
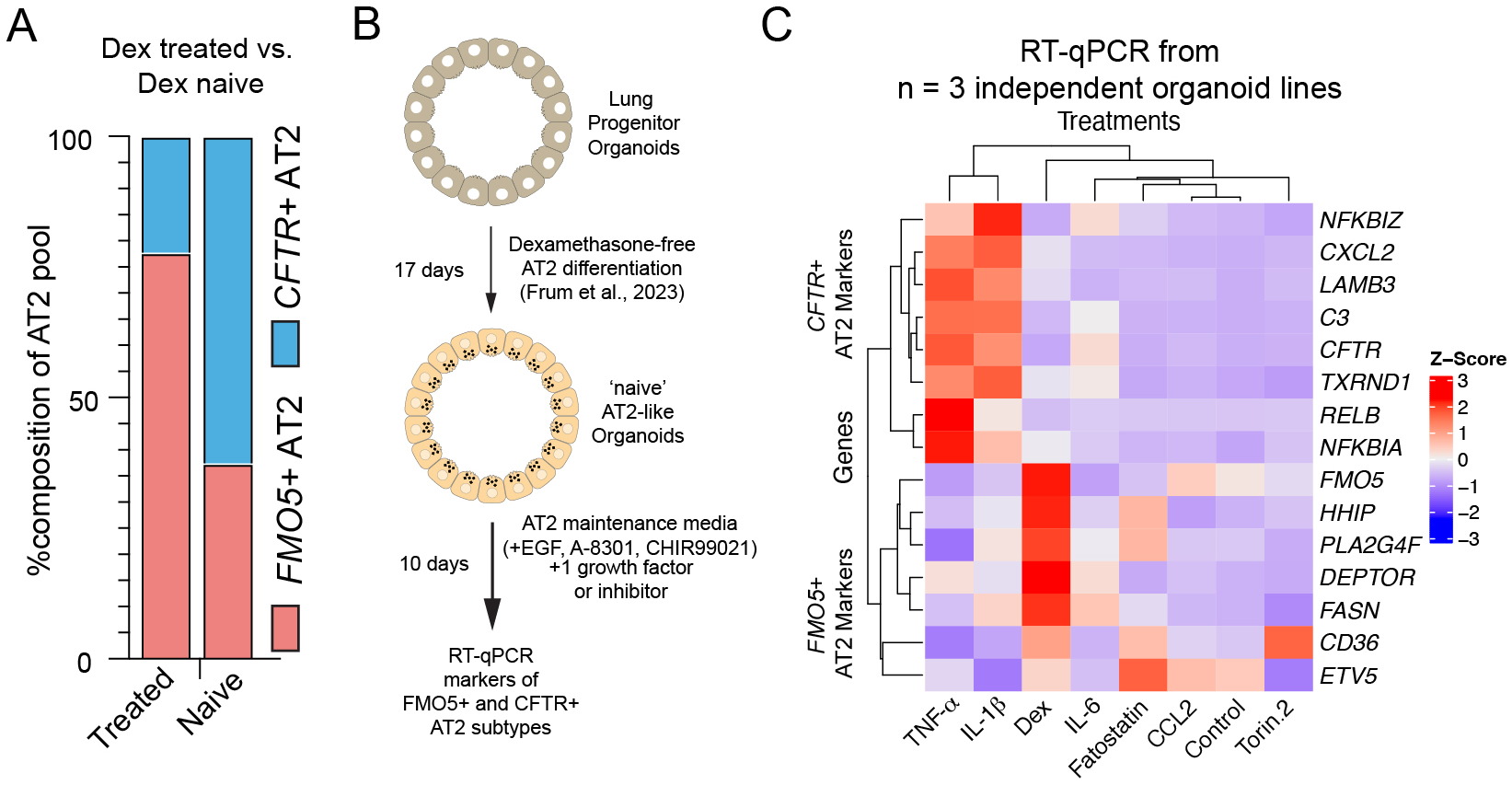
Glucocorticoid and Inflammatory Signals Drive *FMO5+* and *CFTR+* AT2 Transcriptional States In Vitro. (A) Bar graph showing the proportion of *FMO5*+ and *CFTR*+ AT2 cells in early postnatal lung specimens separated by dexamethasone treatment status. Specimens from patients who received dexamethasone contained a higher proportion of *FMO5*+ AT2 cells, whereas dexamethasone-naïve specimens contained a higher proportion of *CFTR*+ AT2 cells. (B) Experimental design. Human lung progenitor organoids were differentiated for 17 days toward AT2-like organoids under dexamethasone-free conditions^58^, then cultured for an additional 10 days in AT2 maintenance media supplemented with one cytokine, growth factor, or small-molecule inhibitor. RT-qPCR was performed to quantify marker genes of *FMO5*+ and *CFTR*+ AT2 transcriptional states. (C) Heatmap showing Z-scored RT-qPCR expression values for markers of *FMO5*+ and *CFTR*+ AT2 transcriptional states across treatments (n = 3 independent organoid lines). *FMO5*+ and *CFTR*+ AT2 markers segregated as expected based on their mutually exclusive expression in tissue. Dexamethasone treatment upregulated FMO5+ AT2 markers, while IL-1β and TNF-α induced *CFTR*+ AT2 markers.

Given that TNF-α and IL-1β are known drivers of inflammation and are pro-regenerative in mice^61–63^, whereas Dex is a potent anti-inflammatory steroid, our results suggest that the inflammatory environment shapes the balance of AT2 cell states in the early postnatal lung.

### Delayed maturation across cell types is a common feature of early postnatal lung disease

To apply our early postnatal lung cell atlas toward a better understanding of cellular alterations in neonatal lung disease, we interrogated diseases associated with the developing lung. This included four patients with respiratory distress in the setting of prematurity (Table S6). Two patients had a clinical diagnosis of evolving bronchopulmonary dysplasia (BPD) with early organizing acute lung injury on pathology and two patients had a clinical diagnosis of severe BPD^11^, with severe deficient alveolarization and pulmonary arterial hypertension on pathology; one of severe BPD patients also had interstitial fibrosis characteristic of “old BPD”^64^. These cases will be termed ‘evolving BPD’ and ‘organized BPD’ in the remainder of the manuscript. Notably only one of these patients had received dexamethasone (Table S6). We also examined five individuals with pulmonary interstitial glycogenosis (P.I.G.) (Table S7), an idiopathic disorder frequently associated with altered alveolar growth and/or acute neonatal lung injury in which glycogen-laden mesenchymal cells accumulate in the interstitial regions of alveoli^65,66^. To facilitate comparisons across datasets, we first performed label transfer from the healthy postnatal atlas to the disease specimens, merging epithelial subclusters (*FMO5*+ and *CFTR*+ AT2 to AT2; AT1-c1, AT1-c2, AT1-c3 to AT1) to generate a common set of annotations across all specimens that are agnostic to transcriptional heterogeneity in the early postnatal alveolar epithelium (Fig. 5A-B).

**Figure 5.**
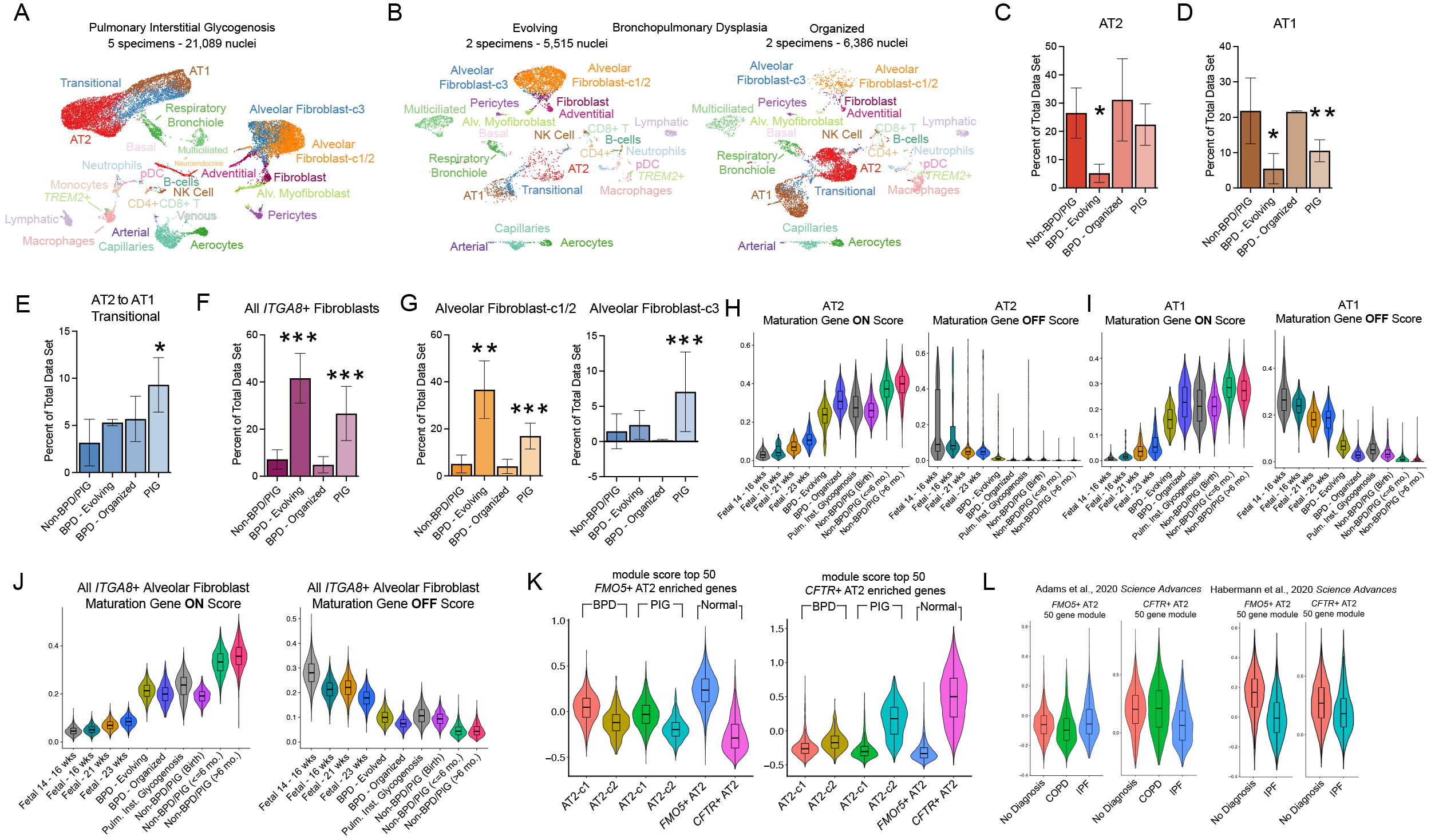
Persistent Mesenchymal Dominance, Failure of Postnatal Maturation, and Repression of CFTR+ AT2 Signatures Are Hallmarks of Early Postnatal Lung Diseases. (A) UMAP of integrated snRNA-seq data from five specimens diagnosed with pulmonary interstitial glycogenosis (P.I.G.). Cell type identities were assigned by label transfer from the histologically normal early postnatal reference atlas. (B) UMAP of integrated snRNA-seq data from four specimens diagnosed with bronchopulmonary dysplasia (BPD), with cell identities transferred from the non-diseased early postnatal atlas. The UMAP is split to contrast specimens with evolving versus organized pathology based on histologic classification. (C) Bar chart comparing the percentage of AT2 cells in non-diagnosed, P.I.G., and BPD specimens. *p < 0.05 relative to non-diagnosed samples by Wilcoxon t-test treating each specimen as a replicate. AT2 cells are significantly reduced in evolving BPD. (D) Bar chart comparing the percentage of AT1 cells across specimen categories. *p < 0.05; **p < 0.01 relative to non-diagnosed samples. AT1 cells are underrepresented in evolving BPD and P.I.G. (E) Bar chart comparing the percentage of AT2 to AT1 transitional cells across specimen categories. *p < 0.05 relative to non-diagnosed samples. Transitional cells are increased in P.I.G. (F) Bar chart showing the proportion of all I*TGA8*+ alveolar fibroblast-c1/2 cells across specimen categories. ***p < 0.001 relative to non-diagnosed samples. Evolving BPD and P.I.G. exhibit excessive ITGA8+ fibroblast abundance. (G) Bar chart showing the relative abundance of alveolar fibroblast subclusters (CCL2-negative, left; CCL2-positive, right) across specimen categories. **p < 0.01; ***p < 0.001 relative to non-diagnosed samples. Excess alveolar fibroblast cells in evolving BPD are predominantly alveolar fibroblast c1/2 cells, whereas P.I.G. contains both alveolar fibroblast-c1/2 and alveolar fibroblast-c3 cells. Similar to evolving BPD, organized BPD also lacks alveolar fibroblast-c3 cells. (H–J) Violin plots showing module scores for maturation-associated gene programs. The “maturation-on” score reflects the top 100 genes enriched in early postnatal relative to fetal cells, while the “maturation-off” score reflects the top 100 genes enriched in fetal relative to postnatal cells. Shown for (H) AT2, (I) AT1, and (J) ITGA8+ fibroblasts, aligned across datasets by label transfer. Disease samples exhibit impaired activation of maturation-on programs and retention of fetal-like programs across many cell types. (K) Violin plot showing module scores for *FMO5*+ and *CFTR*+ AT2 gene sets across bifurcated AT2 clusters from BPD, P.I.G., and non-diagnosed specimens. The *CFTR*+ AT2 signatures is present in P.I.G. but not in BPD. (L) Violin plots showing module scores for *FMO5*+ and *CFTR*+ AT2 gene sets in AT2 clusters from chronic obstructive pulmonary disease and idiopathic pulmonary fibrosis datasets^28,84^. Both datasets show consistent repression of the CFTR+ AT2 gene program in IPF.

Following alignment of cell type identities by label transfer, we compared cell composition across datasets (Fig. 5C-H, Fig. S5A). Evolving BPD showed the most profound disruption of cell type composition in the epithelium, with marked depletion of AT2 and AT1 cells (Fig. 5C-D). P.I.G. retained normal AT2 levels but had increased AT2 to AT1 transitional cells and fewer AT1 cells, suggesting an impaired AT2 to AT1 transition or maintenance of differentiated AT1 cells (Fig. 5C-E). Organized BPD showed no significant changes in epithelial cell number or cell type composition (Fig. 5C-E). Evolving BPD and P.I.G. had greatly elevated representation of AF cells (defined as all *ITGA8*+ cells, Fig. 5F). Strikingly, within the pool of AFs, both types of BPD contained very few AF-c3 mapping cells and were dominated by cells with mapping to an AF-c1/2 (Fig,. 5G, Fig. S5B), while in contrast P.I.G. contained robust populations of both AF-c1/2 and AF-c3 cells (Fig. 5G, Fig. S5B). Although significant alterations in cell composition were detected in the immune and endothelial compartments, the overall size of the difference was small compared to differences in the epithelium and mesenchyme (Fig. S5A, Fig. 5C-F).

These findings highlight persistent mesenchymal dominance as a common pathological axis underlying evolving BPD and P.I.G. In both diseases, excess mesenchymal cells map to an AF identity. Pathology becomes divergent when considering AF subcluster identity, with both evolving and organized BPD lacking AF-c3 cells. In the epithelium, P.I.G. contains an excess of transitional cells, while AT2 and AT1 cells are significantly reduced in evolving BPD. In contrast, epithelial cell type composition appears normal in organized BPD (Fig. 5C-E, Fig. S5A).

Given that alveoli in our organized BPD patients have normal cell type composition, but patients have severe disease with poor lung function, we hypothesized that the cells failed to properly mature in these lungs. To assess the maturation status of each cell type in diseased specimens, we developed cell type-specific gene sets that capture developmental transitions using recently published fetal data (14 to 16 weeks) and additional 16-, 21-, 23-week post-conception samples sequenced for this study (Table S8) and our early postnatal lung data (stratified as ‘at birth’ (n = 2, <1 week), ≤6 months old (n = 10), >6 months old (n = 11). To identify these gene sets, we first performed label transfer to map fetal cell identities to our early postnatal atlas (Fig. S5C).

We noted that label transfer identified a minority of distally localized fetal mesenchymal cells as AF-c1/2 or AF-c3, instead assigning them to the intermediate fibroblast cluster located between myofibroblasts, adventitial, and AF cells in the early postnatal cell atlas (Fig. S1A). Therefore, we also generated an additional cell subset of all *ITGA8*+ mesenchymal cells across specimens and time points. For each cell type, we then identified a ‘maturation gene set’ by comparing gene expression between fetal and early postnatal cell types, defining genes that are upregulated (“on”) (Table S9) or downregulated (“off”) (Table S10) during normal maturation. To benchmark this approach, we also aligned additional fetal specimens (16, 21, and 23 weeks) to the postnatal atlas (Fig. S5D). We then scored (see Methods) all fetal, early postnatal, and disease-state cells for enrichment of “on” and “off” gene sets for each cell type. This approach accurately recapitulated developmental progression in the normal specimens, with cells from 16-, 21– and 23-weeks, and the time-binned early postnatal specimens scoring in a stepwise fashion for upregulated and downregulated maturation gene sets across windows of early postnatal development (Fig. 5H-J; Fig. S5E-O). We note that many immune cell types did not follow this pattern (Fig. S5P-V), likely due to their context-dependent development.

This approach highlighted dramatic post birth maturation in almost all cell types in the normal specimens, particularly for ‘on’ scores, representing genes that are upregulated during development (Fig. 5H-J, Fig. S5E-V). In contrast, post birth maturation across most cell types in the disease specimens appeared to maintain at near birth levels, indicating an arrest in maturation. In general, this effect was strongest in the evolving BPD specimens, followed by P.I.G. and was particularly evident in AT2 cells (Fig. 5H), AT1 cells (Fig. 5I), aerocytes (Fig. S5E), capillaries (Fig. S5F), and mesenchymal cell types, whether compared on the basis of label transfer (Fig. S5K-O) or on the basis of shared *ITGA8* expression (Fig. 5J). The few cells in evolving BPD specimens that mapped to an AF-c3 identity scored much higher than at birth for the ‘off’ score, indicating a specific failure to shut down fetal gene expression programs (Fig. S5L), while organized BPD was distinguished from others by more severe delays in the onset of maturation genes in alveolar myofibroblasts (Fig. S5K). All disease specimens contained a population of AT1 cells with extremely low ‘on’ maturation scores, emphasizing that defects in AT1 maturation are a common feature of neonatal lung disease that may limit gas-exchange function (Fig. 5I).

### *FMO5*+ and *CFTR*+ AT2 transcriptional signatures are modified in lung disease

We were interested in the observation that AF-c3 cells are absent in BPD specimens, given that AF-c3 cells are associated with *CFTR+* AT2s in healthy lungs (Fig. 3). We therefore examined *FMO5+* and *CFTR+* AT2 states in disease specimens by extracting and subclustering AT2s at low resolution to yield two AT2 subclusters per disease cohort. Disease AT2 subclusters were then scored for *FMO5+* and *CFTR+* AT2 transcriptional signatures. This analysis showed that *FMO5+* to *CFTR+* AT2 transcriptional axis was muted in BPD (Fig. 5K). Moreover, BPD was distinguished from P.I.G. by the specific lack of cells scoring high for a *CFTR+* AT2 signature (Fig. 5K). We confirmed that the *CFTR*+ AT2 signature was also diminished in BPD samples relative to normal in an additional cohort of publicly available samples^34^ (Fig. S5W). In light of the fact that P.I.G. is frequently a transient process in the absence of significant comorbidities^67^, we speculate that a lack of *CFTR+* AT2s and AF-c3 cells is a pathologic feature of BPD.

Based on our observation that *CFTR+* AT2 cells are lost in some lung diseases, we expanded our analysis to chronic lung diseases in adults by examining published data sets containing cohorts of chronic obstructive pulmonary disease (COPD) and idiopathic pulmonary fibrosis (IPF) patients^28,29^. As expected, based on our earlier analysis (Fig. S2H) we were unable to find a robust *FMO5+* and *CFTR+* axis of transcriptional heterogeneity in the adult specimens. However, by comparing AT2s from disease specimens to the *FMO5+* and *CFTR+* AT2 signatures, we observed a reduction in the *CFTR+* AT2 signature across IPF datasets, with the *FMO5+* AT2 signature also depressed in one of the datasets (Fig. 5L). The reduction of *CFTR+* and *FMO5+* transcriptional signatures within the AT2s of diseased lungs suggests that maintenance of these transcriptional signatures may be associated with lung health in adults.

## DISCUSSION

The development of the lung’s gas-exchange surface after birth is a critical process for lifelong respiratory health, yet is poorly understood due to limited access of biological specimens at this early stage of life. Our study bridges this knowledge gap by providing the first high-resolution cellular atlas of the healthy human lung from birth to two years of age, with a significant number of histologically confirmed normal biological specimens (n = 23). This resource allowed us to uncover a previously unknown axis of cellular heterogeneity in the alveolar epithelium, fundamentally altering our understanding of AT2 cell biology. We identified two distinct and mutually exclusive AT2 cell states, a lipogenic *FMO5*+ state and a pro-inflammatory/pro-regenerative *CFTR*+ state, that exist in unique cellular niches and are dynamically regulated by the inflammatory environment. These findings provide a new framework for understanding postnatal lung development, regeneration, and the cellular basis of neonatal and adult lung diseases.

A central discovery of our work is that the AT2 lineage is unexpectedly transcriptionally diverse during early postnatal lung development. We hypothesize that early postnatal AT2 states represent a division of labor essential for the dual challenges of postnatal life: ensuring surfactant homeostasis (the *FMO5*+ state) and managing the rapid growth and intermittent remodeling cycles of the developing lung (the *CFTR*+ state). The *FMO5*+ state is characterized by the enrichment of genes involved in lipid metabolism, suggesting these cells are specialized for synthesis of pulmonary surfactant. In contrast, the *CFTR*+ state is defined by an inflammatory gene signature. It is also interesting to note that *CFTR*+ AT2s are distinguished by *ICAM1*, which has been associated with lung morphogenesis and regeneration in mice and human^55,68^. Our functional experiments confirm this, showing that the *CFTR*+ state is induced by TNFα and IL-1β, key cytokines that play a role in alveolar regeneration in mice and primates^61–63,69^. Additionally, our spatial transcriptomic analysis shows that *CFTR*+ AT2s associate with specific cluster of AF cells (AF-c3) that share gene signatures with pro-regenerative alveolar fibroblast populations in mice^53–55^. We also show that AF-c3s are a source of SEMA3C, which has recently been hypothesized to be a critical cue for normal lung function that is absent in BPD^34^. Together, these findings suggest that the *CFTR*+ AT2 population is indicative of a localized pro-regenerative state and may be a key source of the lung’s innate capacity for regeneration.

Our work also demonstrates that the balance between the two AT2 states is not fixed but is instead exquisitely sensitive to the surrounding inflammatory milieu, with profound clinical implications. We found that dexamethasone, a glucocorticoid widely used in neonatal care to promote lung maturation and diminish lung inflammation^70–74^, potently induces the *FMO5*+ lipogenic state *in vitro* and is associated with a striking predominance of this state in treated patients. While this shift likely confers a short-term benefit to lung function by boosting surfactant production, our results would argue that it comes at the cost of depleting the pro-regenerative *CFTR*+ AT2 pool. This finding provides a mechanistic explanation for the long-observed trade-offs of steroid therapy, which are thought to promote AT2 maturation, encourage surfactant production, and reduce lung inflammation. Notably, steroid therapy has not led to a decrease in the number of BPD cases^75–79^. It is also interesting to note that the *CFTR*+ AT2 state was associated with an NF-κB gene signature and may therefore be important for proper long-term alveolar development or maturation, as suggested by studies on NF-κB signaling in mice^80–82^. Therapies that suppress the *CFTR*+ AT2 state could inadvertently compromise the lung’s developmental trajectory and its capacity for future repair.

This new understanding of AT2 biology sheds light on the implications of neonatal lung disease states, which may persist across the human lifespan. In our analysis of BPD and P.I.G., we observed a profound arrest of cellular maturation across multiple lineages, indicating a failure to execute the normal postnatal developmental program. Strikingly, we found that BPD is characterized by a near-complete absence of the AF-c3 cells and their associated *CFTR*+ AT2 population. This loss of the pro-regenerative AT2 niche appears to be a common pathological feature, as we also observed a significant repression of the *CFTR*+ AT2 signature in other BPD datasets, and adult lung datasets such as IPF and COPD. This suggests that the inability to mount or sustain this regenerative program may be a fundamental driver of chronic lung disease, regardless of age. The *CFTR*+ AT2 state and its associated niche may therefore represent a novel therapeutic target and a valuable biomarker for monitoring lung health and disease progression.

In conclusion, our work charts the cellular landscape of the human lung during a critical and dynamic period of postnatal development. By integrating single-cell and spatial transcriptomics with functional organoid models, we have defined two functionally specialized alveolar niches, uncovered novel transcriptional states in the AT2 lineage, and identified relevant signaling cues that place *CFTR*+ and *FMO5*+ AT2 transcriptional states under the control of an inflammatory-reparative axis. These findings not only redefine our understanding of the alveolar stem cell compartment but also provide a powerful new lens through which to view lung health, regeneration, and the pathogenesis of both pediatric and adult respiratory diseases.

## MATERIALS AND EXPERIMENTAL METHODS

### Experimental Model and Subject Details

#### Human lung specimens

All research utilizing human lung specimens, (14 – 23 weeks post-conception lung, birth to 2 years early postnatal lung) was performed with the approval of the University of Michigan Institutional Review Board. Human lung tissue section from specimens pre-birth were processed by the University of Washington Laboratory of Developmental Biology and were presumably normal. All specimens are de-identified. Tissue was processed within 24 hours of isolation and transported in Belzer-UW Cold Solution (ThermoFisher, Cat#NC0952695) at 4°C. Early postnatal lung specimens were obtained from either autopsies or surgical resections (Table S1; Table S6-8) which were split a matched pieces were either snapped frozen for snRNA-seq processing or archived for histology, immunostaining and spatial transcriptomics by formalin fixation and paraffin embedding.

### BTP organoid establishment, maintenance and differentiation to AT2-like cells

Primary BTP organoid cultures from 15 – 18.5 weeks post conception lung tissue were established and maintained as previously described (Miller, 2020). Matrigel was diluted to 8 mg/mL before use. Bud Tip Progenitor Media consisted of DMEM/F-12 (Corning, Cat#10-092-CV), 100U/mL penicillin-streptomycin (ThermoFisher, Cat#15140122), 1X B-27 supplement (ThermoFisher, Cat#17504044), 1X N2 supplement (ThermoFisher, Cat#17502048), 0.05% Bovine Serum Albumin (BSA, Sigma, Cat#A9647), 50µg/mL L-ascorbic acid (Sigma, Cat#A4544), 0.4 µM 1-Thioglycerol (Sigma, Cat#M1753), 50nM all-trans retinoic acid (Sigma, Cat#R2625), 10ng/mL recombinant human FGF7 (R&D Systems, Cat#251-KG), and 3µM CHIR99021 (APExBIO, Cat#A3011). BTP organoids were needle passaged every 7 to 10 days with 10 µM Y-27632 (APExBIO, Cat#A3008) added to the media for the first 24 hours after each passage. After three passages and an additional 3 – 5 days to permit recovery of fragmented organoids, organoids were differentiated towards an AT2 cell identity by removing Bud Tip Progenitor Media and adding CKAB AT2 differentiation media (Frum et al., 2023) consisting of DMEM/F12, 100U/mL penicillin-streptomycin, 1x B-27 supplement, 1x N2 supplement, 0.05% BSA, 50 µg/mL L-ascorbic acid, 0.4 µM 1-Thioglycerol, 10 ng/mL recombinant human FGF7, 3 µM CHIR99021, 100 ng/mL BMP4 (R&D Systems, Cat#314-BPE) and 1 µM A83-01 (APExBIO, Cat#A3133) and passaged after 7 days treatment. After an additional 10 days CKAB AT2 differentiation media was replaced with AT2 expansion media consisting of DMEM/F12, 100U/mL penicillin-streptomycin, 1x B-27 supplement, 1x N2 supplement, 0.05% BSA, 50 µg/mL L-ascorbic acid, 0.4 µM 1-Thioglycerol, 50 ng/mL EGF (R&D Systems Cat#236EG), 1 µM A83-01 and 3 µM CHIR99021) with or without the addition of 1 growth factor, small molecule or metabolite as detailed in Table S4.

### Primary AT2 Organoid Establishment and Maintenance

Primary AT2 cells were purified from dissociated early postnatal lung tissue by magnetic assisted cell sorting using an HTII-280+ antibody as detailed in published protocols (Tata, Nature Protocols, 2020) and cultured as organoids embedded in Matrigel (8 mg/mL) domes under SFFF media (Katsura et al., 2020). Primary AT2 organoids were passaged every 20 – 30 days. After two passages organoids were again sorted by laser assisted flow cytometry for HTII-280+/NGFR-cells to obtain a pure population of HTII-280+ AT2 cells. AT2 cells were passaged by dissociation into single cells using TrypLE Express (ThermoFisher, Cat#12605010), quenched with equal volume DMEM/F12 + 10% FBS (Corning Cat#35015CV) and embedding in Matrigel at 1000 cells/µl. Recombinant hIL-1b (ProteinTech, Cat#HZ-1164) and recombinant TNF-α (ProteinTech, Cat#HZ-1014) were added to SFFF media at a concentration of 100 ng/mL. Dexamethasone (Sigma, Cat#D4902) was added to SFFF media at a concentration of 50 nM.

### Preparation of Tissue for Fluorescence *In Situ* Hybridization, Protein Immunofluorescence and Spatial Transcriptomics

Lung tissue specimens were fixed for 24 hours with 10% Neutral Buffered Formalin at room temperature with gentle rocking. Samples were washed three times with DNase/RNase-Free Water (ThermoFisher, Cat#10977015) and dehydrated through 25%, 50%, 75% and 100% methanol washes followed by washes in 100% and 70% ethanol. Specimens were paraffin processed in an automated tissue processor through 70%, 80%, 2×95%, 3×100% ethanol, 3x xylene and 3x paraffin. All wash steps and parrafin processing steps occurred for 60 minutes at room temperature. Tissue was embedded into paraffin blocks and sectioned at 5 µm thickness using a microtome. Slides were baked for 1 hour at 60°C prior to staining.

### Fluorescence *In Situ* mRNA Hybidization

mRNA hybridization was perfromede using the RNAscope Multiplex Fluorescent V2 assay (ACDBio, Cat#323100) using TSA-Cy3 (Akoya Biosciences Cat#NEL744001KT) and TSA-Cy5 (Akoya Biosciences, Cat# NEL745E001KT) following recommendations provided by the manufacturer. Protease treatment was performed with Protease Reagent IV for 5 minutes. Antigen retrieval was performed for 15 minutes.

### Protein Staining by Indirect Immunofluorescence

For protein Immunofluorescence costaining, slides were washed in 1x Phosphate Buffered Saline (PBS) (Corning, Cat#21040) after final HRP treatment and immediately put into antibody blocking solution (0.1% Tween-20, 5% Normal Donkey Serum, PBS) for 1 hour, followed by treatment with mouse primary antibody against Surfactant Protein C (Santa Cruz Biotechnology, Cat#sc518029) diluted 1 to 200 in antibody blocking solution for 24 hours at 4°C, 3x washes with PBS, treatment with AffiniPure Donkey Anti-Mouse IgG (H + L) conjugated to Alexa Fluor 488 (Jackson ImmunoResearch, Cat#715545151) diluted 1 to 400 and DAPI (Sigma, Cat# D9542) diluted 1 to 1000 in antibody blocking solution for 1 hour at room temperature, 3x washes with PBS. Approximately 10 µl of ProLong Gold (ThermoFisher, Cat#P369300) was applied to tissue sections before mounting with 1.5 thickness glass coverslips.

### Confocal Imaging

Imaging was performed on a Nikon AXR confocal microscope using a 40x Silicon Oil objective at 2x digital zoom and 2048×2048 pixel resolution with z-step size of 0.5 microns. Z-stacks were combined to form maximum projections in Nikon Elements v5.

### Tissue Preparation for Xenium Spatial Transcriptomics Analysis

Xenium slides were removed from –20°C storage and allowed to come to room temperature for 30 minutes and then were placed on a 42°C slide warmed and coated with DNAse/RNAse free water (Corning, Cat# 46000CM). Small sections from multiple specimens were carefully placed within the sample placement area.

Most of the water was removed when sections had completely flattened. Slides dried on the slide warmer for three hours before transport to the Advanced Genomics Core. Xenium slides were processed by the Advanced Genomics Core using the Xenium *In Situ* Gene Expression with Cell Segmentation workflow (10X, Doc#CG000749).

### RNA extraction, Reverse mRNA Transcription and Quantitative Polymerase Chain Reaction (RT-qPCR)

Organoids were recovered from Matrigel by trituration using a P1000 tip and transferred to a 1.5 mL tube where they were pelleted by centrifugation at 300xG for 5 minutes, media and Matrigel were removed and then pellets were flash frozen in liquid nitrogen at stored at –80°C. RNA was isolated from organoid pellets using the MagMax-96 Total RNA Isolation kit (ThermoFisher, Cat#AM1830). RNA yield and quality was determined on a Nanodrop 2000 sprectrophotometer. Reverse Transcription using the SuperScript VILO cDNA kit (Invitrogen) was performed in triplicate for each biological replicate for 60 minutes at 42°C using 200 ng RNA input and a reaction volume of 20 µl. cDNA was diluted 1:4 and 1/80^th^ of the final reaction was used for each RT-qPCR measurement. RT-qPCR was performed on a Step One Plus Real-Time PCR System (ThermoFisher, Cat#43765592R) using QuantiTect SYBR Green qPCR Master Mix (Qiagen, Cat#204145) with primers at a concentration of 500 nM and 40 cycles of amplification. Primer specificity was determined by a single peak during the melt-cycle of each run. Sequences for RT-qPCR primers used in this manuscript are in Supplementary Table 7. Arbitrary units of target mRNA abundance were normalized to expression of small ribosomal subunit 18 (18S) and calculated as: **2^(18SCt-TargetCt)^ x 10,000.**

### Submission of samples for single nucleus RNA-sequencing (snRNA-seq)

Samples were stored in LN2 until preparation for snRNA-seq. Small pieces from each sample were shaved of a single region specimen and then minced into small rice grain sized fragments using a No.1 Scapel in a 6 cm dish on dry ice. Samples were transferred to 1.5 mL tubes, dissociated by pestle and then nuclei were purified using the 10X Chromium Nuclei Isolation Kit (10X Genomics, Cat#1000493) following the manufacturer’s recommendations, counted using a Countess Automated Cell Counter (v2) and resuspended at 1000 cells/µl in PBS + 1% BSA (Mitenyl, Cat#). The University of Michigan Advanced Genomics Sequencing Core prepared libraries using the Chromium Next GEM Single Cell 3’ GEM, Library and Gel Bead Kit v3.1 (10X Genomics, Cat#PN1000128) targeting 7500 nuclei per specimen. snRNA-sequencing libraries were sequenced to a projected average read depth of 80,000 reads per nuclei using a NovaSeq 6000 with S4 300 cycle reagents.

### Analysis of snRNA-seq data

#### Ambient RNA Correction

Reads were mapped to human genome (GRCh38-2020-A) and gene expression matrices generated using CellRanger v7. Raw matrices were used as input for CellBender v0.30, which re-called nuclei and corrected for ambient RNA at a false-positive rate of 0.01. CellBender corrected gene expression matrices were imported into Seurat v5 in RStudio v1.4 running R v4.1.

#### Preprocessing/QC Filtering

Only nuclei within the following thresholds were considered for further analysis: between 500 to 7500 features, more than 1000 unique molecules sequenced, less than 5% mitochondrial RNA reads and less than 7.5% ribosomal reads.

#### Data Integration, Dimensional Reduction, Clustering

For each specimen data was normalized using Seurat::NormalizeData() and then all specimens were integrated using a mutual nearest neighbors batch correction implemented by SeuratWrappers::RunFastMNN() in SeuratWrappers v0.3.5 using 2000 features. The first 30 dimensions of the mutual neighbors reduction was used to generate a Uniform Manifold Approximation and Projection (UMAP) by Seurat::RunUMAP(). Nearest-neighbor graph construction was performed using Seurat::FindNeighbors(). Louvain clustering was performed at a resolution of 1.0 using Seurat::FindClusters().

#### Additional QC

Clusters were inspected for mutually exclusive expression of major cell class markers (Epithelial: *CDH1, EPCAM;* Endothelial: *PECAM1*, Immune: *PTPRC*, Mesenchymal: *PDGFRA, PDGFRB)*. Clusters of cells at the center of the UMAP coexpressing markers of multiple major cell classes were removed. We speculate these data points are doublets or ambient RNA. After removal, Data Integration, Dimensional Reduction and Clustering were performed again.

#### Cell Type Annotation

Clusters were first annotated as Epithelium, Mesenchyme, Immune or Endothelium based on unique expression of the major cell class markers identified above (Fig. 1B). Each of these annotations was subclustered, using the following top dimensions of the integrated mutual nearest neighbors reduction calculated by Seurat::RunFastMNN() on the complete dataset as input for Seurat::RunUMAP() and Seurat::FindNeighbors(): Epithelium: 20, Mesenchyme: 25, Immune: 16, Endothelium: 15. Louvain clustering was performed using Seurat::FindClusters() at the following resolutions: Epithelium: 0.8, Mesenchyme: 0.4, Immune: 0.3, Endothelium: 0.15. At this point clusters were annotated to minor cell classes based on known markers (i.e. Airway vs Alveolar, Lymphoid vs. Myeloid, etc) (Fig. 1C, D). Some of these minor cell classes were further subclustered to achieve a cell type level annotation (Vessels, Lymphoid, Myeloid), while all others were annotated on the cluster structure evident at the first round of subclustering. Cell type annotations (Fig. 1E) were consistent with known markers (Fig. 1H) of cell type identity.

#### Differential Gene Expression

Genes significantly enriched in each cluster were calculated by Wilcoxon Rank Sum test using Seurat::FindAllMarkers() considering all genes expressed in at least 25% of cluster cells and a log fold expression value greater than 0.25 and excluding all mitochondrial or ribosomal genes. Genes significantly differentially expressed between two clusters were calculated by Wilcoxon Rank Sum test using Seurat::FindMarkers() with the same cutoffs and exclusions.

#### Gene Module Scoring

Gene set modules were scored in Seurat v5 using the gene set method developed by Tirosh et al.,. Brifely, each module gene is scored against a set of 100 control genes that are selected from a bin of genes with similar expression level. The expression level of genes in the module set relative to control genes is then calculated. Gene sets for the *FMO5*+ and *CFTR*+ AT2 transcriptional signatures were determined by direct comparison of both AT2 cell subtypes using Seurat::FindMarkers() and ranked by log fold expression. Gene sets for the canonical AT2 and signaling AT2 defined from sequencing adult lung were generated from the top 50 enriched genes by direct comparison of canonical and signaling AT2s using Seurat::FindMarkers() on dataset ID:syn21560510 downloaded from http://www.synapse.org/#!Synapse:syn21041850. Gene sets for Aerocytes and Capillary Endothelial cells were taken from Supplemental Table 5 of Schupp et al., 2021^38^.

#### Label Transfer

For label transfer annotated reference data was re-processed and integrated using principle component analysis approach. Each individual dataset was first normalized by NormalizeData(), a set of 2000 variable features defined by FindVariableFeatures() were scaled using ScaleData() followed by principle component determination using RunPCA(). Data was then integrated on anchors determined by Seurat::FindIntegrationAnchors() using Seurat::IntegrateData() with 30 dimensions. The resulting ‘integrated’ assay was scaled by Seurat::ScaleData() and Principal Components (PCs) for the ‘integrated’ assay were determined by Seurat::RunPCA(). Query datasets were processed the same way. Anchors for label transfer were determined using Seurat::FindTransferAnchors() with k.filter = 200 using the ‘integrated’ assays of both the reference and query data and 30 dimensions. Cell type annotations of ‘labels’ were transferred from reference top query data by Seurat::TransferData() using 30 dimensions.

#### Cell Communication Analysis

Putative signaling interactions between cell types were determined using CellChat v2 using the default cell signaling database database and without the use of the provided protein-protein interaction database. Only cell signaling pathways annotated as “Secreted Signaling” were analyzed and interactions comprising less than 10 cells were removed. For Fig. S3A-F this was performed on two CellChat objects sharing all AT1 and Transitional subclusters and including either AT2 subtype and it’s associated fibroblast population (i.e. *FMO5*+ AT2s and Alveolar Fibroblast-c1/2 or *CFTR*+ AT2s and Alveolar Fibroblast-c3), excluding other mesenchymal, immune and endothelial populations. Both datasets were merged into a single CellChat object for comparison. For Fig. S3G,H separate CellChat objects were generated by splitting snRNA-seq data into data from individuals enriched for the *FMO5*+ AT2s or individuals enriched for *CFTR*+ AT2s. These datasets contained all cell types and were merged into a single CellChat object for comparison.

#### Trajectory Inference

For visualization of lineage geometry the embedding was computed with Seurat::RunUMAP(n.components = 3), yielding a three-dimensional manifold. The resulting coordinates were used as the input space for Slingshot trajectory inference and for 3-D visualization using the plotly or rgl packages in R. Trajectories were inferred with Slingshot^83^. Trajectory roots were specified as the two AT2 subclusters and endpoints as the AT1-c1 subclusters; AT2 to AT1 transitional cells were left unspecified to allow data-driven path fitting. Slingshot fitted principal curves and computed lineage-specific pseudotime for each cell. To visualize branch geometry, we rendered the first three embedding dimensions in 3D and overlaid curve segments

### Xenium Data Analysis

#### Preprocessing/QC Filtering

Centroids and Segmentation coordinates and Gene Expression counts were determined by Xenium Onboard Analysis v4.0 and imported into R using Seurat::ReadXenium(). Gene Expression counts were converted to a Seurat object using Seurat::CreateSeuratObject(). Coordinates for centroids and segmentations were first converted into a field of view using Seurat::CreateFOV() and then appended to the Seurat object. Segmentations with less than 25 gene expression counts were excluded from the analysis.

#### Label Transfer

To align low-complexity 480 probe Xenium data with higher complexity snRNA-seq data the reference data was transformed using Seurat::SCTransform() with 3000 variable features. Each specimen was processed individually, also undergoing SCTransformation using 250 variable features. Any Xenium probes expressed in over 95% of cells were excluded from analysis. Anchors between each specimen and the snRNA-seq reference were calculated using FindTransferAnchors() using the SCT assay of both datasets, 20 dimensions, k.filter = 200, and considering only the variable features from the Xenium specimen. Cell type annotations from the snRNA-seq data were then transferred to the Xenium specimen using TransferData(), with anchors weighted by the PCs of the Xenium specimen.

#### Merged Embedding of Xenium Specimen Data

All xenium specimens were merged into a single object and then transformed by Seurat::SCTransform(). SCTransformed values were used as input for RunPCA(). UMAP was determined by Seurat::RunUMAP() using 30 principle components. For Fig. 3B metadata containing identities inferred by label transfer was transferred from each individual xenium specimen to the merged embedding of all xenium specimen data.

#### Niche Analysis

For Fig. 3C-E each individual specimen underwent ‘niche’ analysis using Seurat::BuildNicheAssay() with niches.k = 4 and neighbors.k = 20 using cell type identities inferred by label transfer from snRNA-seq data (Fig. 3B). The cell type composition of each niche in each specimen was merged into a data table. Pearson’s correlation between cell type compositions of each niche identified in this analysis was generated using stats::cor() and this was used as input for pheatmap::pheatmap() with clustering of columns and rows. Niches of similar composition were clustered together and then mapped back onto sections to correlate them with anatomical regions in each specimen (i.e. interstitium, bronchiole, alveolar) (Fig. 3D). The composition of the two identified alveolar niches was extracted and analyzed in Excel (Microsoft) with Prism (Graphpad) used for plotting (Fig. 3E). For Fig. S3F-I each specimen underwent ‘niche’ analysis using Seurat::BuildNicheAssay() with niches.k = 2 and neighbors.k = 10 using cell type identities determined by hand annotation of AT2 and Alveolar Fibroblast populations (Fig. S3B-D). Pearson’s correlation was performed and used as input for pheatmap::pheatmap with clustering of columns and rows. The composition of the two alveolar niches was analyzed in Excel (Microsoft) with plotted with Prism (Graphpad).

#### Distance Analysis

Using hand annotation of AT2 and Alveolar Fibroblast subclusters (Fig. S3B-D) we calculated a symmetric mean nearest-neighbor distance using coordinates extracted by Seurat::GetTissueCoordinates(): the average of distances from each cell of type A to the closest cell of type B neighbor and from cell each cell of type B to the closest cell of type A as determined by RANN::get.knnx(). Distances were computed for all sections and aggregated into a single table. Significant differences between cell-type pair distances were determined using a one-way ANOVA (base::aov()) followed by Tukey’s Honest Significant Difference test for pairwaise comparisons (base::TukeyHSD).

#### Join Count Test

To examine the spatial association of *FMO5*+ and *CFTR*+ AT2s we extracted coordinates of hand annotated *FMO5*+ and *CFTR*+ AT2s for each section and offsetting y to generate a field of all five specimens. A distance-band neighbor graph was constructed using spded::dnearneigh. Spatial association was tested by joincount.test() and the reported Z value and two-sided p-values for *FMO5*+ to *FMO5*+ AT2 and *CFTR*+ to *CFTR*+ AT2 self-association at that distance are reported in Fig. S3 using distances from 10 to 30 microns in 1 micron increments.

## CODE AVAILABILITY

Code used for analysis of single nuclei transcriptomic and spatial transcriptomic data is available from: http://github.com/jason-spence-lab/Frum-et-al.-2025a.git

## DATA AVAILABILITY

Fully processed and annotated RObjects for all single nuclei transcriptomic datasets analyzed in this manuscript are available at Zenodo (https://zenodo.org/uploads/17361944). Xenium Ranger output from spatial transcriptomic analysis is also available at Zenodo (https://zenodo.org/uploads/17155546). At the time of publication in a peer-reviewed journal all raw sequencing read data will be available at EMBL-ArrayExpress and is available by request to.

## Supporting information

Table S1 – Sample MetaData For Histologically Normal Early Postnatal Specimens

Table S2 – Gene Signatures of FMO5+ and CFTR+ AT2s in Early Postnatal Lung

Table S3 – Gene Signatures of Canonical and Signaling AT2s in Adult

Table S4 – Xenium 480 Custom Panel Targets

Table S5 – Compounds Tested on Primary AT2 Organoids

Table S6 – Sample MetaData For Lung Specimens Diagnosed With Bronchopulmonary Dysplasia

Table S7 – Sample MetaData For Lung Specimens Diagnosed With Pulmonary Interstitial Glycogenosis

Table S8 – Sample MetaData For 16-, 21-, 23-week Lung Specimens

Table S9 – Maturation "ON" gene sets by Cell Type

Table S10 – Maturation "OFF" gene sets by Cell Type

Table S11 – Primers for RT-qPCR

## ACKNOWLEDGEMENTS

We express gratitude to the patients who donated tissue samples to make this research possible. This work was supported by the Chan Zuckerberg Initiative (CZI) Seed Network (2019-002440) to J.R.S. and CZI Pediatric Network (2021-237566), an advised fund of the Silicon Valley Community Foundation, to G.H.D. and J.R.S., and the National Heart, Lung, and Blood Institute (NHLBI; R01HL119215 and R01HL166139) to J.R.S. S.G.C. was supported by the National Science Foundation (NSF; DGE2241144). P.P.H. was supported by the Judith Tam ALK Lung Cancer Research Initiative, the Rogel Cancer Center Fellowship, and the National Cancer Institute (K08CA28675). I.A.G. and the Washington Laboratory of Developmental Biology was supported by Eunice Kennedy Shriver National Institute of Child Health & Human Development (NICHD; R24HD000836).

## FIGURE LEGENDS

**Figure S1.**
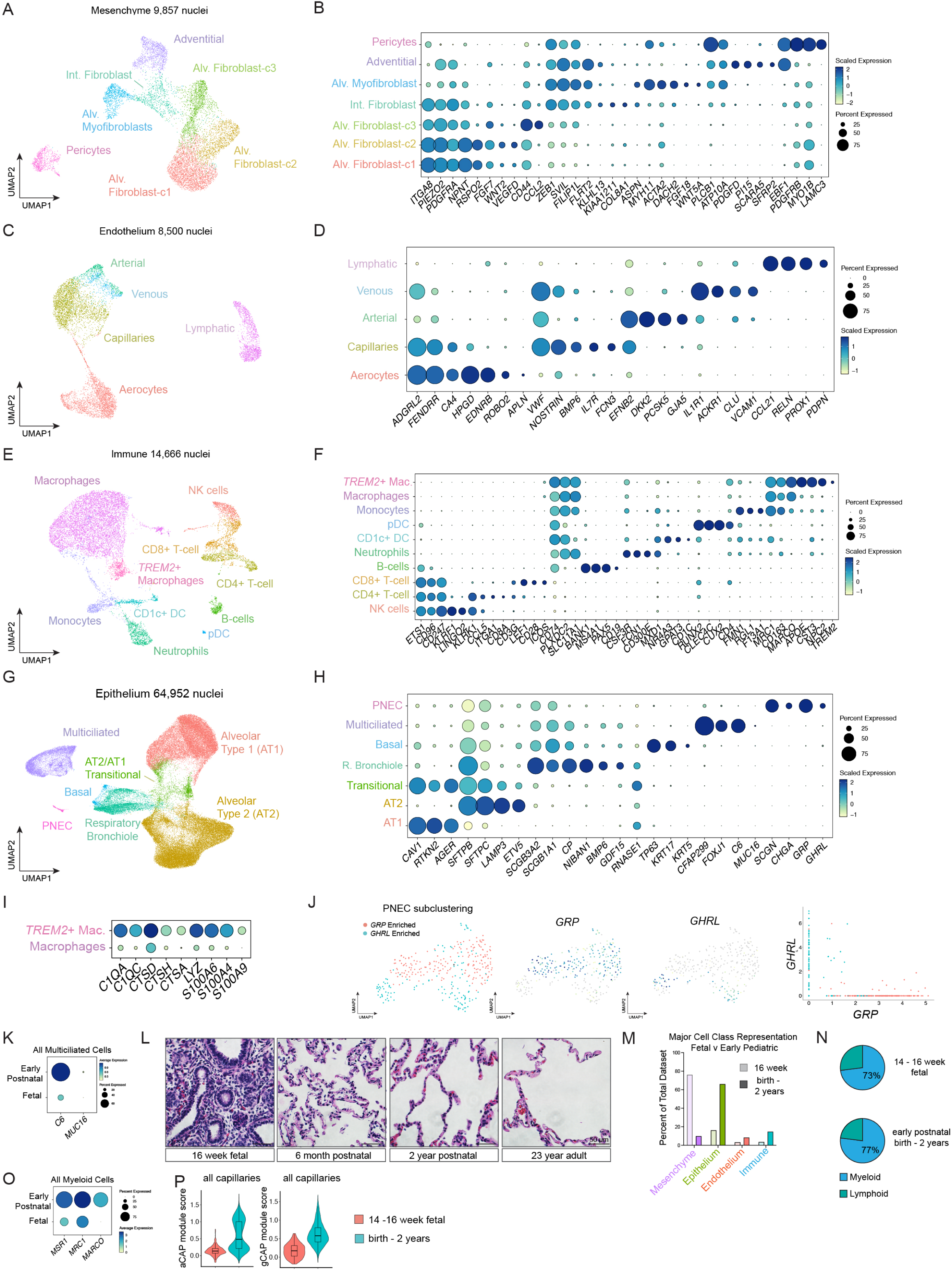
Subclustering of Major Cell Classes and Identification of Known Cell Types in the Early Postnatal Lung. (A) UMAP embedding of mesenchymal subclusters, annotated based on enrichment of known marker genes shown in (B). (B) Dot plot illustrating shared and distinguishing markers of alveolar fibroblast-c1, –c2, and –c3, subclusters, adventitial fibroblasts, alveolar myofibroblasts, pericytes, and shared marker expression across intermediate fibroblasts. (C) UMAP embedding of endothelial subclusters with annotations based on known marker enrichment highlighted in (D). (D) Dot plot showing markers distinguishing capillary aerocytes and capillary endothelial cells, and distinguishing markers among larger-vessel endothelial populations (venous, arterial, lymphatics). (E) UMAP embedding of immune subclusters with annotations based on marker enrichment shown in (F). (F) Dot plot illustrating shared and distinguishing markers across lymphoid and myeloid populations. (G) UMAP embedding of epithelial subclusters annotated by marker expression shown in (H). (H) Dot plot showing markers distinguishing epithelial populations including AT1, AT2, AT2 to AT1 transitional, basal, multiciliated, respiratory bronchiole, and pulmonary neuroendocrine (PNEC) cells. (I) Dot plot showing genes enriched in *TREM2*+ macrophages indicative of an activated or polarized state. (J) UMAP of PNEC subclusters enriched for *GRP* or *GHRL*, with feature and scatter plots demonstrating mutually exclusive expression of subset markers in the early postnatal distal lung. (K) Dot plot showing that multiciliated cells exhibit a *C6*+/*MUC16*+ transcriptional profile in the early postnatal distal lung. (L) Hematoxylin-and-eosin staining of fetal (14–16 weeks), early postnatal (birth to two years), and adult alveoli. Scale bar = 50 μm. (M) Comparison of cellular composition between 14–16-week fetal and early postnatal distal lung by major cell class, highlighting the developmental shift from mesenchymal to epithelial dominance. (N) Relative myeloid and lymphoid contributions to the immune compartment are similar between fetal and early postnatal specimens. (O) Dot plot comparing macrophage marker expression between fetal and early postnatal lungs reveals emergence of *MARCO+* resident macrophages postnatally. (P) Violin plot comparing aCAP (aerocyte) and gCAP (capillary) module scores, derived from Travaglini et al. 2020, across all capillary cells in fetal and early postnatal datasets, demonstrating the emergence of specialized capillary signatures, particularly aCAP/aerocyte enrichment, after birth.

**Figure S2.**
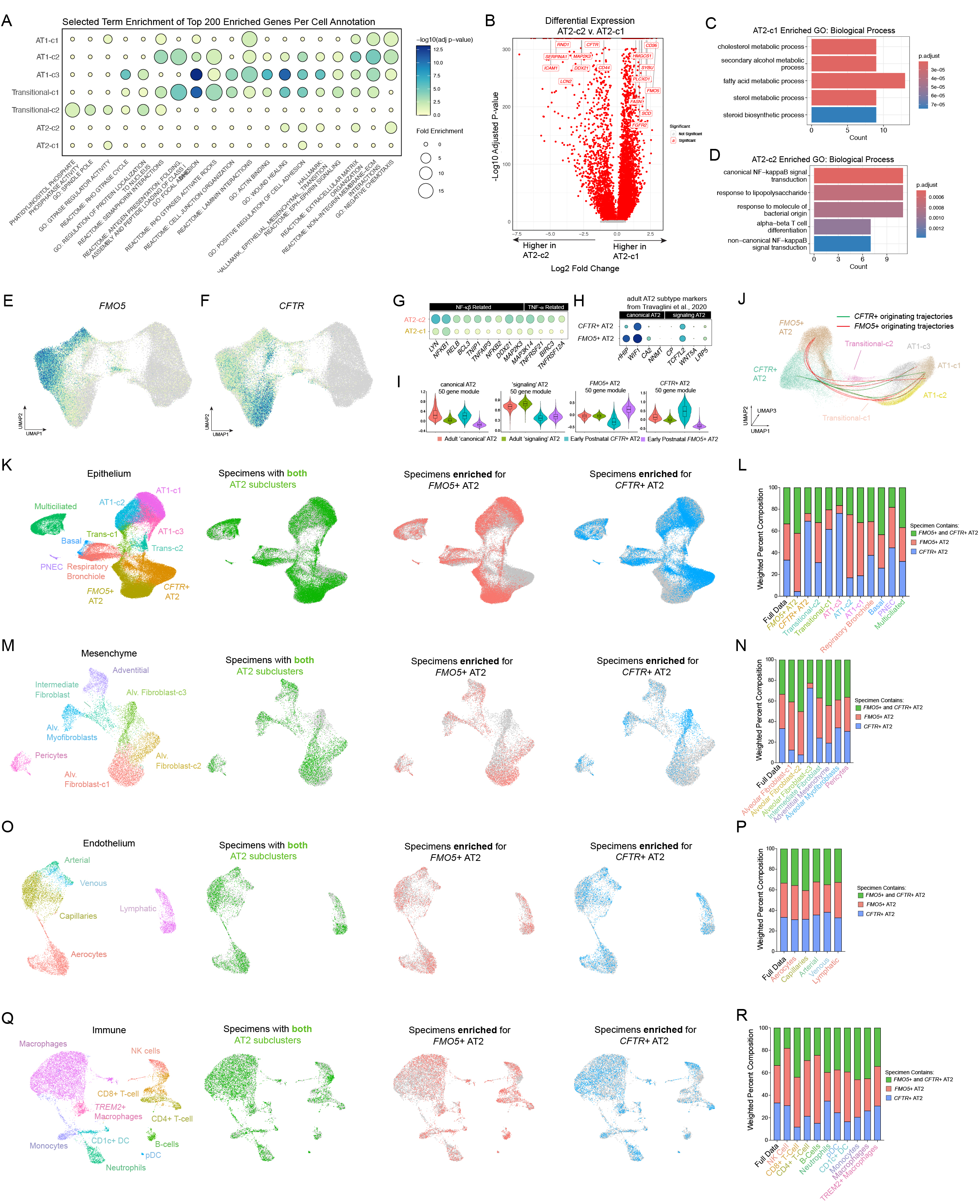
Functional Divergence and Cell Type Associations of AT2 Subclusters in the Early Postnatal Lung. (A) Dot plot of gene set enrichment analysis showing biological processes and pathways enriched across alveolar epithelial subclusters, emphasizing terms enriched in AT2 to AT1 transitional and AT1 clusters. (B) Volcano plot showing significantly differentially expressed genes between AT2 subclusters. (C) The top five enriched GO Biological Process terms for AT2-c1 markers converge on lipid metabolic processes. (D) The top five enriched GO Biological Process terms for AT2-c2 markers converge on immune-responsive pathways, including NF-κB and other immune related signaling. (E) Feature plot showing the spatial distribution of the AT2-c1 marker *FMO5*. (F) Feature plot showing the spatial distribution of the AT2-c2 marker *CFTR*. (G) Dot plot showing enrichment of known targets of NF-κB and TNF-α signaling in AT2-c2, supporting an immune-responsive phenotype in AT2-c2 cells. (H) Dot plot showing expression of adult AT2 subtype markers^27^ within early postnatal *FMO5*+ and *CFTR*+ AT2 populations. (I) Gene-module scoring comparing early postnatal and adult datasets. Left: scoring of early postnatal AT2 subclusters using 50-gene modules defining canonical and signaling AT2 subtypes in the adult lung. Right: scoring of adult AT2 subclusters using 50-gene modules derived from early postnatal *FMO5*+ and *CFTR*+ AT2 transcriptional axes. (J) Slingshot trajectory analysis on a three-dimensional UMAP of alveolar epithelial cells reveals differentiation trajectories originating from *FMO5*+ and *CFTR*+ AT2 subtypes and converging on AT1 cells through AT2 to AT1 transitional intermediates. (K) Epithelial UMAP showing annotations and overlaying cells from specimens classified as having comparable proportions of both AT2 subtypes (green), enriched for AT2-c1 (red), or enriched for AT2-c2 (blue). (L) Weighted composition of epithelial cell types across specimen classifications, adjusted so that each category contributes equally to the overall epithelial compartment (left column, “Full Data”). (M) Mesenchymal UMAP showing annotations and overlaying cells from the same specimen categories (green = both AT2 subtypes, red = AT2-c1 enriched, blue = AT2-c2 enriched). (N) Weighted composition of mesenchymal cell types across specimen classifications, normalized as in (L). (O) Endothelial UMAP showing annotations and specimen overlays as in (M). (P) Weighted composition of endothelial cell types across specimen classifications, normalized as in (L). (Q) UMAP of immune cell subclustering showing annotations and specimen overlays as in (M). (R) Weighted composition of immune cell types across specimen classifications, normalized as in (L).

**Figure S3.**
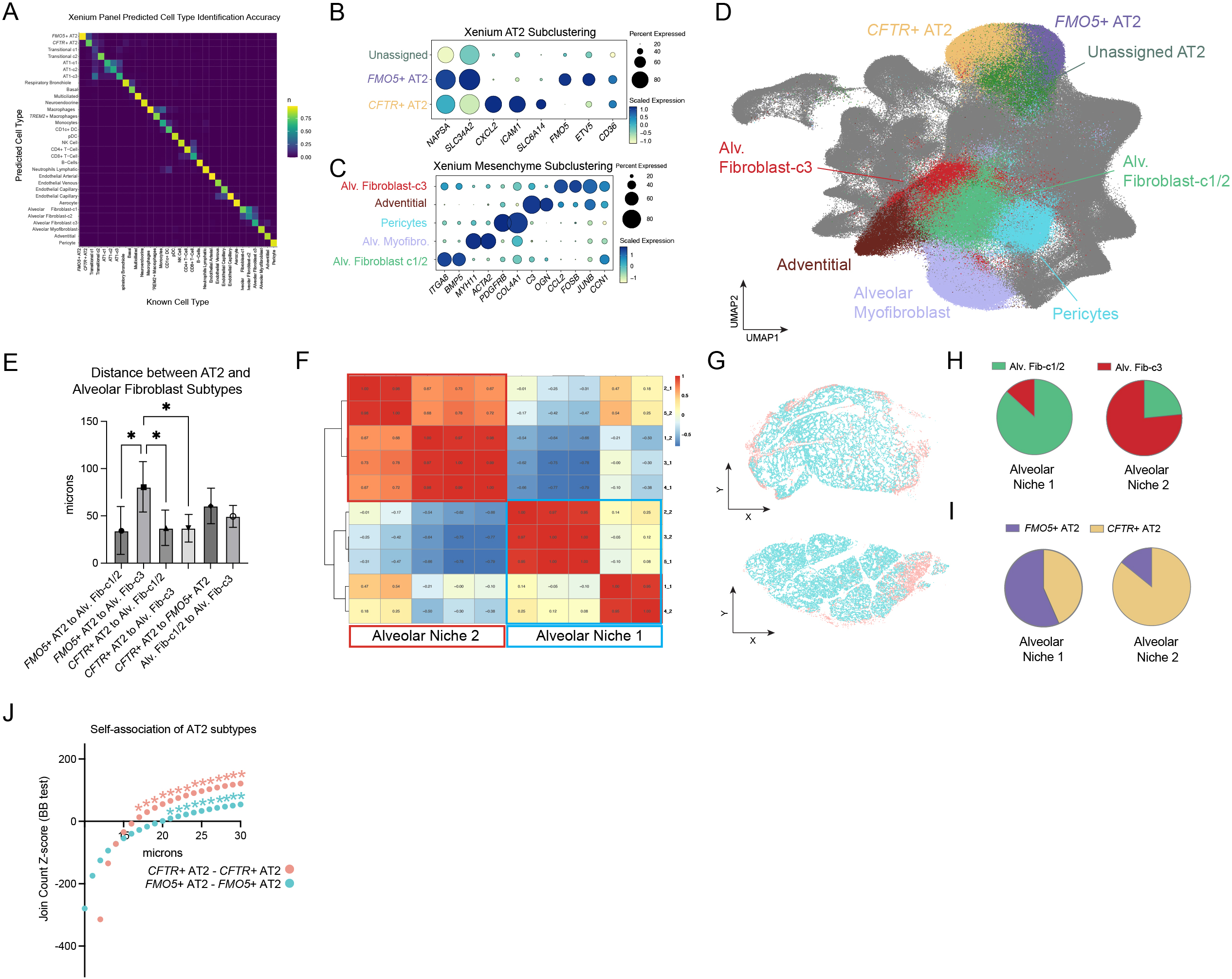
*FMO5*+ and *CFTR*+ AT2s are Spatially Associated with Distinct Alveolar Fibroblast Subtypes. (A) Confusion matrix showing the accuracy of cell type identification using the custom 480-gene Xenium probe set (y-axis), validated against cell type identities from early postnatal snRNA-seq data (x-axis). (B) Dot plot showing marker gene expression for manually annotated AT2 cells in the Xenium dataset. This approach identifies a subset of unassigned AT2 cells lacking FMO5 and CFTR expression, which were excluded from analyses in panels E–J. (C) Dot plot showing marker gene expression for manually annotated mesenchymal cells in the Xenium dataset, distinguishing alveolar fibroblast-c3 from alveolar fibroblast-c1/2 populations. (D) Integrated UMAP of all segmented spatial transcriptomic cells, colored by manually annotated cell type identities. (E) Quantification of the average spatial distance between AT2 and alveolar fibroblast subtypes across tissue sections. FMO5+ AT2 cells are positioned farther from alveolar fibroblast-c3 cells than from alveolar fibroblast-c1/2 cells, whereas CFTR+ AT2 cells are closely associated with alveolar fibroblast-c3 cells. (F) Heatmap showing Pearson’s correlation of niche cell composition derived from Seurat v5’s k-means–based niche analysis using only manually annotated AT2 and mesenchymal cells. Two distinct alveolar niche clusters emerge, corresponding to the FMO5+ and CFTR+ AT2 microenvironments. (G) Spatial maps of two early postnatal lung sections colored by niche classification based on (F). (H) Pie charts showing that alveolar niche 1 is enriched for and alveolar fibroblast-c1/2 cells, while alveolar niche 2 is enriched for alveolar fibroblast-c3 cells. (I) Pie charts showing that alveolar niche 1 is enriched for *FMO5+* AT2 cells and *CFTR+* AT2 cells. (J) Join count spatial autocorrelation analysis of AT2 subtype self-association. Z-scores were calculated across increasing spatial distances. Asterisks indicate significant positive association (*p < 0.05). CFTR+ AT2 cells show stronger local self-association than FMO5+ AT2 cells, indicating tighter spatial clustering.

**Figure S4.**
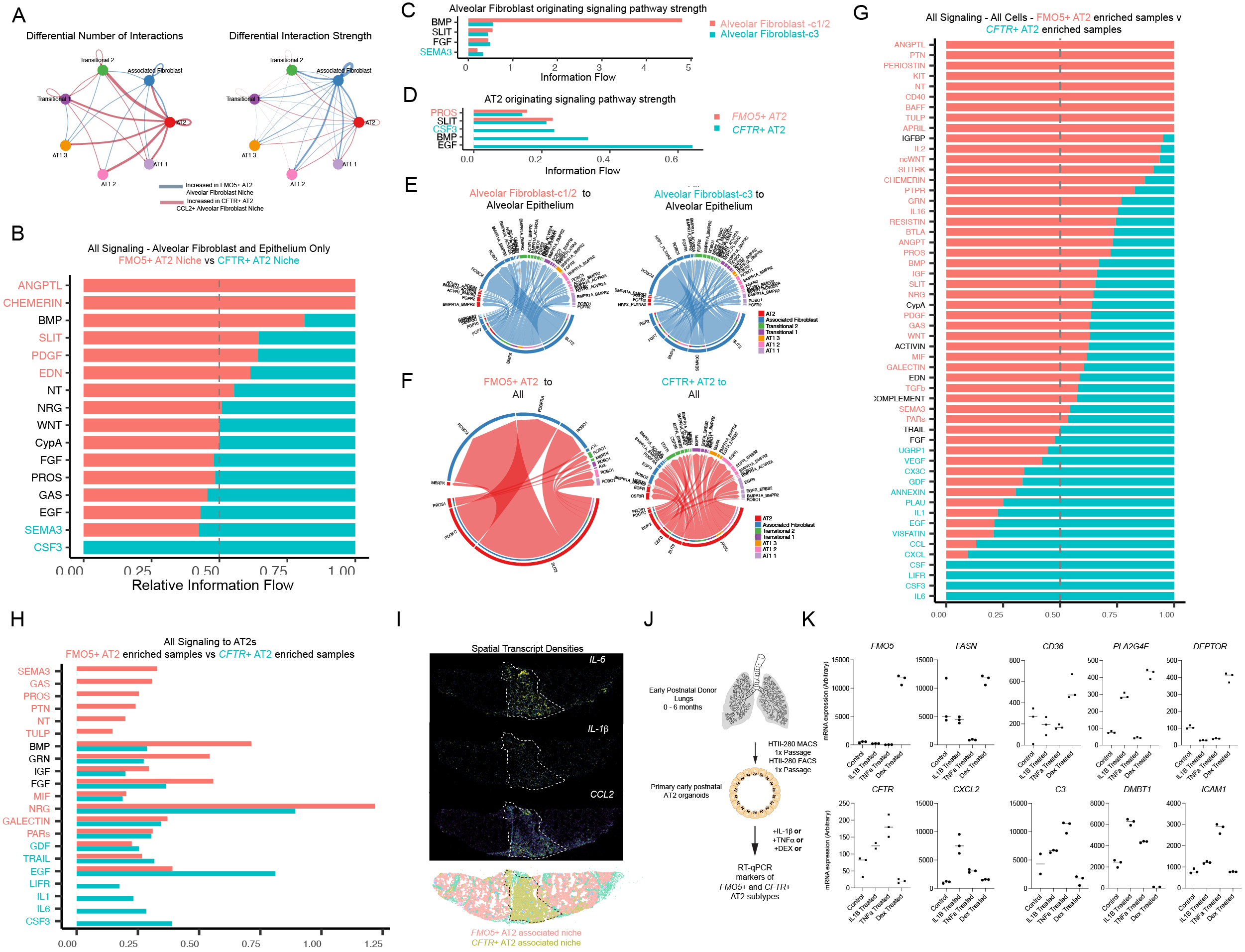
CellChat Analysis of *FMO5*+ and *CFTR*+ AT2 alveolar niches and recapitulation of *FMO5*+ and *CFTR*+ AT2 States in Primary Early Postnatal Organoids. (A–F) CellChat analysis of alveolar epithelial and mesenchymal interactions within *FMO5*+ and *CFTR*+ AT2 niches. The *FMO5*+ AT2 niche includes *FMO5*+ AT2 cells, alveolar fibroblast-c1/2 populations, and all AT2 to AT1 transitional and AT1 subclusters. The *CFTR*+ AT2 niche includes *CFTR*+ AT2 cells, alveolar fibroblast-c3 populations, and the same transitional and AT1 subclusters as the *FMO5+* AT2 niche. AT2 subclusters and their associated fibroblast subsets (i.e. *CFTR*+ AT2 and AF-c3) are referred to as “AT2” and “Associated Fibroblast” in the plots. (A) Differential number of interactions and interaction strength between alveolar epithelial and mesenchymal cell types in each niche. (B) Comparison of pathway-level signaling activity between *FMO5*+ and *CFTR*+ AT2 niches. Pathways highlighted in red are significantly more active in the *FMO5*+ AT2 niche, whereas those in blue are more active in the *CFTR*+ AT2 niche. (C) Comparison of absolute outgoing signaling from alveolar fibroblast-c1/2 cells in the *FMO5*+ AT2 niche and alveolar fibroblast-c3 cells in the *CFTR*+ AT2 niche. Pathways highlighted in blue are significantly more active when alveolar fibroblast-c3 present. (D) Comparison of absolute outgoing signaling from *FMO5*+ and *CFTR*+ AT2 cells within their respective niches. Pathways highlighted in red are more active in the *FMO5*+ AT2 niche; those in blue are more active in the *CFTR*+ AT2 niche. (E) Circle plot showing all ligand–receptor interactions originating from alveolar fibroblast-c1/2 (left) or alveolar fibroblast-c3 cells (right). Circle size indicates total signaling strength from the source fibroblast; segment size indicates the relative contribution of each ligand or receptor. alveolar fibroblast-c3 cells express more *FGF2* and *SEMA3C* relative to alveolar fibroblast-c1/2 cells. (F) Circle plot showing all ligand–receptor interactions originating from *FMO5*+ AT2 (left) or CFTR+ AT2 (right) subclusters. *CFTR*+ AT2 cells uniquely express *CSF3* and *AREG*. (G–H) CellChat comparison between specimens enriched for *FMO5*+ or *CFTR*+ AT2 cells, considering all cell types. (G) Relative pathway-level signaling activity across all cell types. Pathways in red are more active in the *FMO5*+ AT2 niche, and those in blue are more active in the *CFTR*+ AT2 niche. (H) Comparison of total incoming signaling strength to AT2 cells between *FMO5*+ AT2– and C*FTR*+ AT2-enriched specimens. (I) Spatial transcript density maps of *IL-6*, *IL-1*β, and *CCL2* transcripts from Xenium spatial transcriptomics of early postnatal lung sections. Transcripts encoding these ligands are enriched in regions corresponding to the *CFTR*+ AT2-enriched alveolar niche (gold, black dashed outlines). (J) Experimental scheme for generating early postnatal AT2 organoids from donor lungs and directing them toward *FMO5*+ or *CFTR*+ transcriptional states. Briefly, donor lungs (0–6 months) were dissociated to single cells, AT2s isolated via HTII-280 magnetic-assisted cell sorting, and cultured in AT2 organoid media^85,86^ for 30 days. Following one passage and purification by flow-assisted cell sorting for HTII-280+/NGFR-cells, organoids were expended and treated for 10 days with dexamethasone, IL-1β, or TNF-α, then collected for RT-qPCR. (K) RT-qPCR analysis of *FMO5*+ (top panels) and *CFTR*+ (bottom panels) AT2 marker genes. Primary early postnatal AT2 organoids recapitulate *in vivo* AT2 cell states, with IL-1β, or TNF-α promoting *CFTR*+ AT2 marker expression and dexamethasone inducing *FMO5*+ AT2 marker expression.

**Figure S5.**
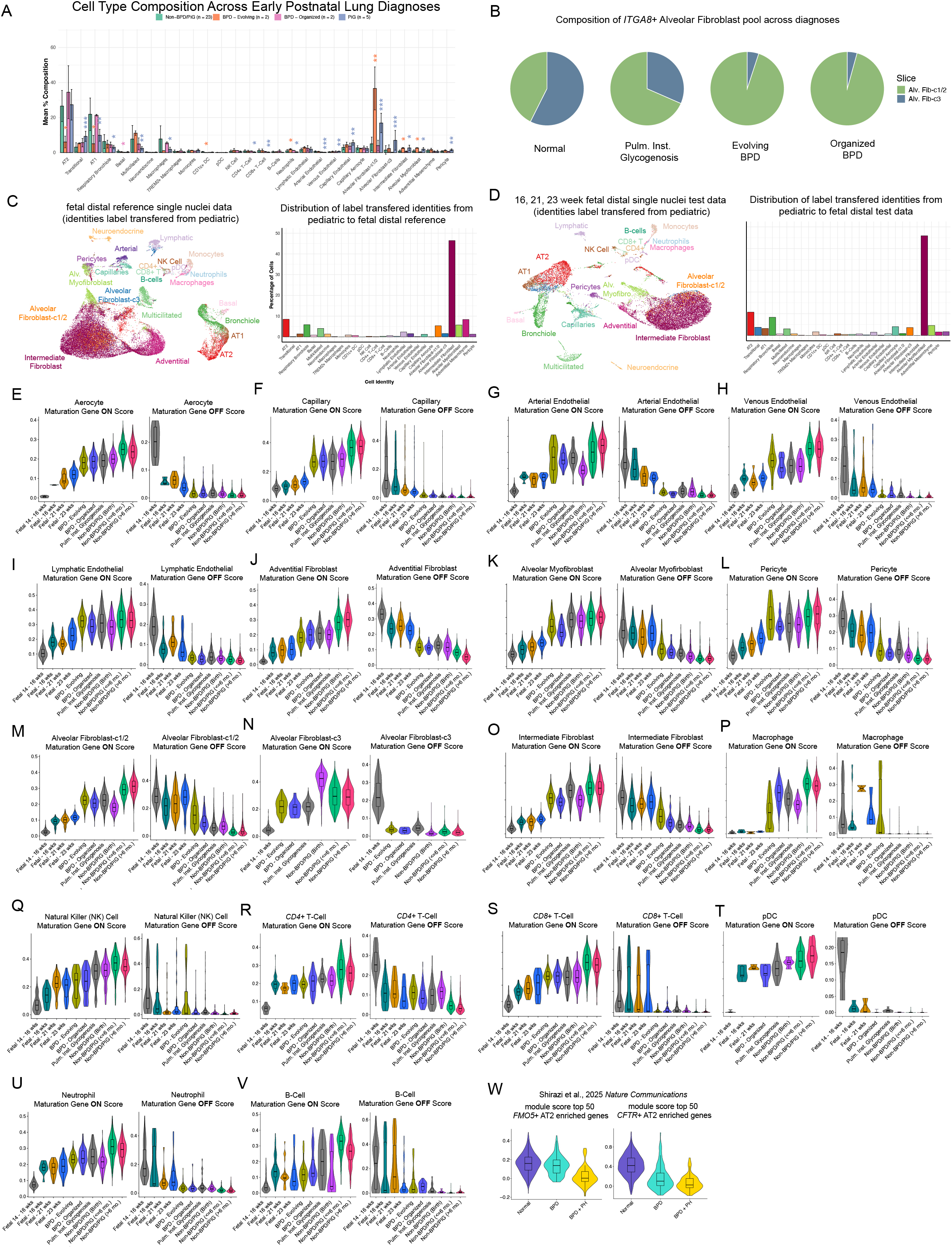
Cell Type Composition and Maturation Scoring Differences Between Fetal, Non-Diagnosed, and Diseased Early Postnatal Lung Specimens. (A) Bar chart comparing mean cell type composition across all non-diagnosed and diagnosed early postnatal lung specimens. Error bars indicate standard deviation. *p < 0.05, **p < 0.01, ***p < 0.001 as determined by Wilcoxon t-test treating each specimen as a biological replicate. (B) Pie charts showing the proportion of mesenchymal cells mapping to alveolar fibroblast-c1/c2 and alveolar fibroblast-c3 populations in non-diagnosed, pulmonary interstitial glycogenosis (P.I.G.), evolving bronchopulmonary dysplasia (BPD), and organized BPD specimens. (C) UMAP of 14 to 16-week fetal distal lung specimens used as a reference to define maturation gene sets, with cell type identities transferred computationally from the early postnatal snRNA-seq atlas (left). Bar chart showing the inferred cell type composition of 14 to 16-week fetal distal samples (right). (D) UMAP of additional 16-, 21-, and 23-week fetal distal lung specimens used as benchmarks for maturation scoring. Identities were transferred computationally from the early postnatal lung atlas (left), and the corresponding inferred cell type composition is shown (right). (E–V) Violin plots showing module scores for maturation gene programs across major cell types. The “maturation-on” score represents the top 100 genes enriched in early postnatal relative to fetal cells, and the “maturation-off” score represents the top 100 genes enriched in fetal relative to postnatal cells. Panels show scores for: (E) aerocytes, (F) capillary endothelial, (G) arterial endothelial, (H) venous endothelial, (I) lymphatic endothelial, (J) adventitial fibroblasts, (K) Alveolar Myofibroblasts, (L) pericytes, (M) alveolar fibroblast-c1/c2 cells, (N) alveolar fibroblast-c3 cells, (O) intermediate fibroblasts, (P) macrophages, (Q) natural killer, (R) CD4+ T Cells, (S) CD8+ T Cells, (T) plasmacytoid dendritic cells (pDCs), U) Neutrophils, and V) B Cells. (W) Violin plots showing module scores for *FMO5*+ and *CFTR*+ AT2 transcriptional signatures in an independent dataset containing BPD and BPD with pulmonary hypertension (BPD + PH) specimens^34^. This data shows the *CFTR*+ AT2 state is reduced in BPD.

